# Vocalization Patterns in Laying Hens - An Analysis of Stress-Induced Audio Responses

**DOI:** 10.1101/2023.12.26.573338

**Authors:** Suresh Neethirajan

## Abstract

This study leverages Convolutional Neural Networks (CNN) and Mel Frequency Cepstral Coefficients (MFCC) to analyze the vocalization patterns of laying hens, focusing on their responses to both visual (umbrella opening) and auditory (dog barking) stressors at different ages. The aim is to understand how these diverse stressors, along with the hens’ age and the timing of stress application, affect their vocal behavior. Utilizing a comprehensive dataset of chicken vocal recordings, both from stress-exposed and control groups, the research enables a detailed comparative analysis of vocal responses to varied environmental stimuli. A significant outcome of this study is the distinct vocal patterns exhibited by younger chickens compared to older ones, suggesting developmental variations in stress response. This finding contributes to a deeper understanding of poultry welfare, demonstrating the potential of non-invasive vocalization analysis for early stress detection and aligning with ethical livestock management practices. The CNN model’s ability to distinguish between pre- and post-stress vocalizations highlights the substantial impact of stressor application on chicken vocal behavior. This study not only sheds light on the nuanced interactions between stress stimuli and animal behavior but also marks a significant advancement in smart farming. It paves the way for real-time welfare assessments and more informed decision-making in poultry management. Looking forward, the study suggests avenues for longitudinal research on chronic stress and the application of these methodologies across different species and farming contexts. Ultimately, this research represents a pivotal step in integrating technology with animal welfare, offering a promising approach to transforming welfare assessments in animal husbandry.

## 1. Introduction

### 1.1. The Communicative Complexity of Hen Vocalizations

The vocalizations of laying hens represent a sophisticated system of communication that goes beyond mere sound production. These vocalizations are a window into the hens’ internal states, conveying critical information about their physical and psychological well-being [1, 2]. For instance, a hen’s cluck may express contentment or signal to chicks to stay close, while a loud squawk often indicates distress or threat [3, 4]. The subtleties in frequency, pitch, and duration of these sounds are not random but are carefully modulated to convey specific messages.

Researchers and poultry caretakers can glean vital insights from these vocal cues. A sudden change in the vocalization pattern could indicate health issues such as disease or injury, or it might reflect environmental stressors like temperature fluctuations or inadequate nutrition [5]. Recognizing these cues promptly can lead to timely interventions, enhancing the overall welfare of the flock.

### 1.2. Social Dynamics and Vocalizations

Within a flock, vocalizations play a pivotal role in social structuring and group cohesion. Hens utilize specific calls to assert dominance, establish social hierarchies, or mediate conflicts, which are essential for maintaining order and reducing aggression within the group [6, 7]. These vocal signals also facilitate cooperative behaviors, such as foraging and nesting [8]. Understanding the nuances of these social communications is crucial for managing flocks effectively, ensuring that social stressors do not compromise the birds’ welfare.

In addition, maternal calls are a critical component of the hen-chick relationship [9]. These sounds play a significant role in the survival of chicks, as they learn to respond to their mother’s calls for food and warning of dangers [10]. Disruptions in these vocal communications can have profound effects on chick development and survival rates.

### 1.3. Vocalizations in Commercial Poultry Production

In the realm of commercial poultry production, monitoring vocal patterns offers a non-intrusive, efficient way to assess the health and welfare of the birds on a large scale. Automated sound analysis systems can detect deviations in vocalization patterns [11], alerting caretakers to potential issues in realtime. This proactive approach can lead to improved flock management strategies, better disease control, and enhanced productivity.

### 1.4. Overview of Stress Impact on Poultry Vocalizations

#### 1.4.1. The Influence of Stress on Vocal Behaviors

The vocal behavior of hens is acutely sensitive to stress. When exposed to stressors, hens often exhibit increased vocalization frequency and altered acoustic properties in their calls [12]. These vocal changes are not just behavioral responses but also reflect underlying physiological stress responses [13]. For example, prolonged stress can impair immune function, reduce growth rates, and even affect egg production [14].

#### 1.4.2. Differentiating Responses to Various Stressors

The type of stressor can elicit distinct vocal responses. Physical stressors, such as temperature extremes or poor living conditions, might result in vocalizations that differ in frequency or duration from those caused by social stressors like overcrowding or pecking order disputes [15]. These differences are crucial for pinpointing the specific stress factors affecting the flock. Understanding how different stressors impact vocal behavior is vital for designing environments and management practices that minimize stress. For instance, modifying housing conditions or adjusting flock densities can significantly reduce stress-induced vocalizations, indicating an improvement in the birds’ well-being.

#### 1.4.3. Chronic Stress and Vocalization Patterns

Chronic stress leads to more profound changes in vocalization patterns [16]. These long-term alterations can be subtle, yet they are indicative of sustained changes in the hens’ physiological and psychological states. Continuous monitoring of these vocal patterns offers a valuable tool for assessing the long-term welfare of the birds. By identifying and addressing the sources of chronic stress, poultry producers can enhance the overall health and productivity of their flocks.

Table 1 contextualizes the study within the broader field of poultry vocalization research and succinctly summarizes recent advancements in analyzing poultry vocal behaviors, offering insights into various methods and key findings that have shaped current understanding. This study builds upon this foundation, employing advanced machine learning techniques to classify and analyze the vocalization patterns of laying hens under stress.

**Table 1.**
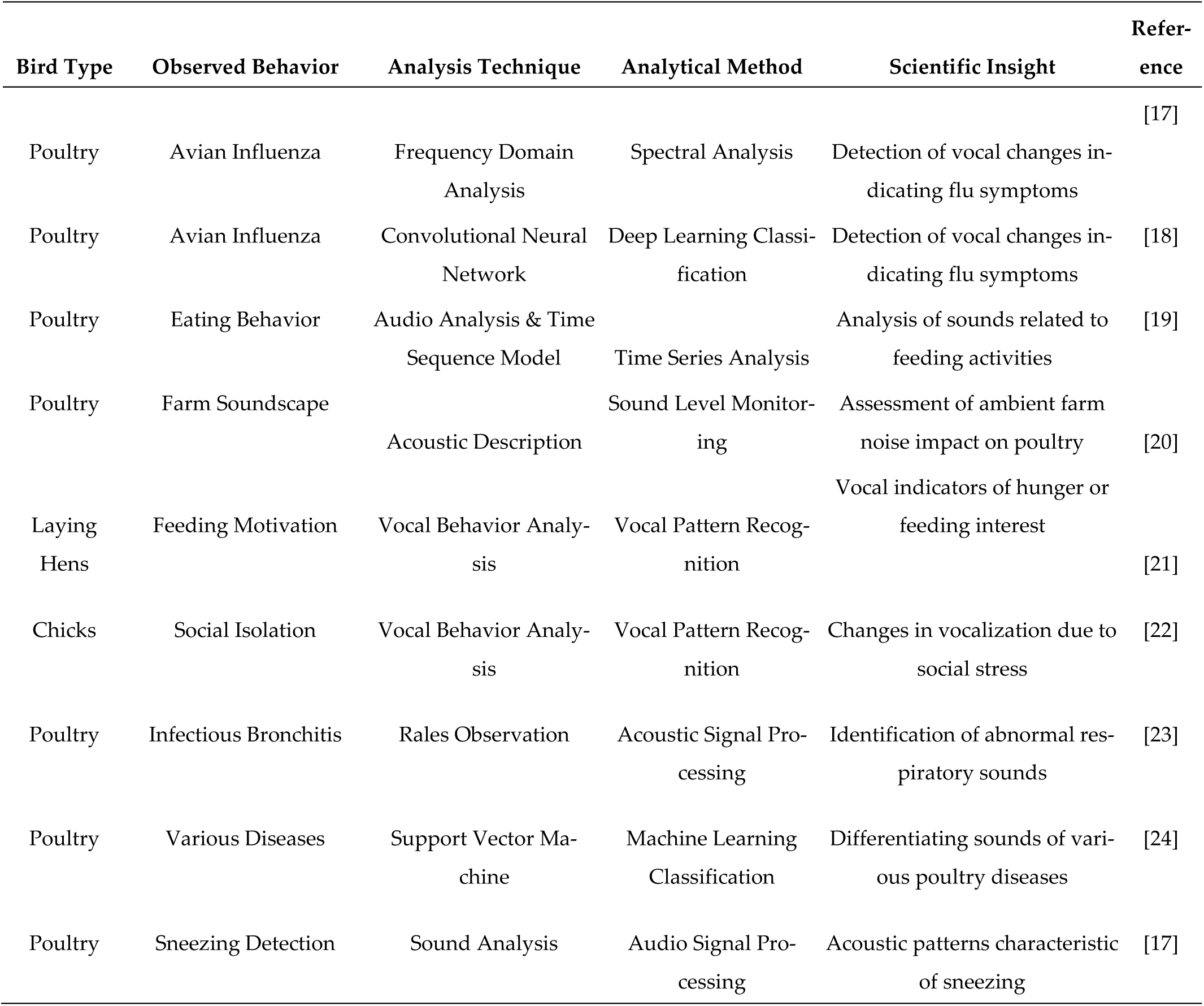
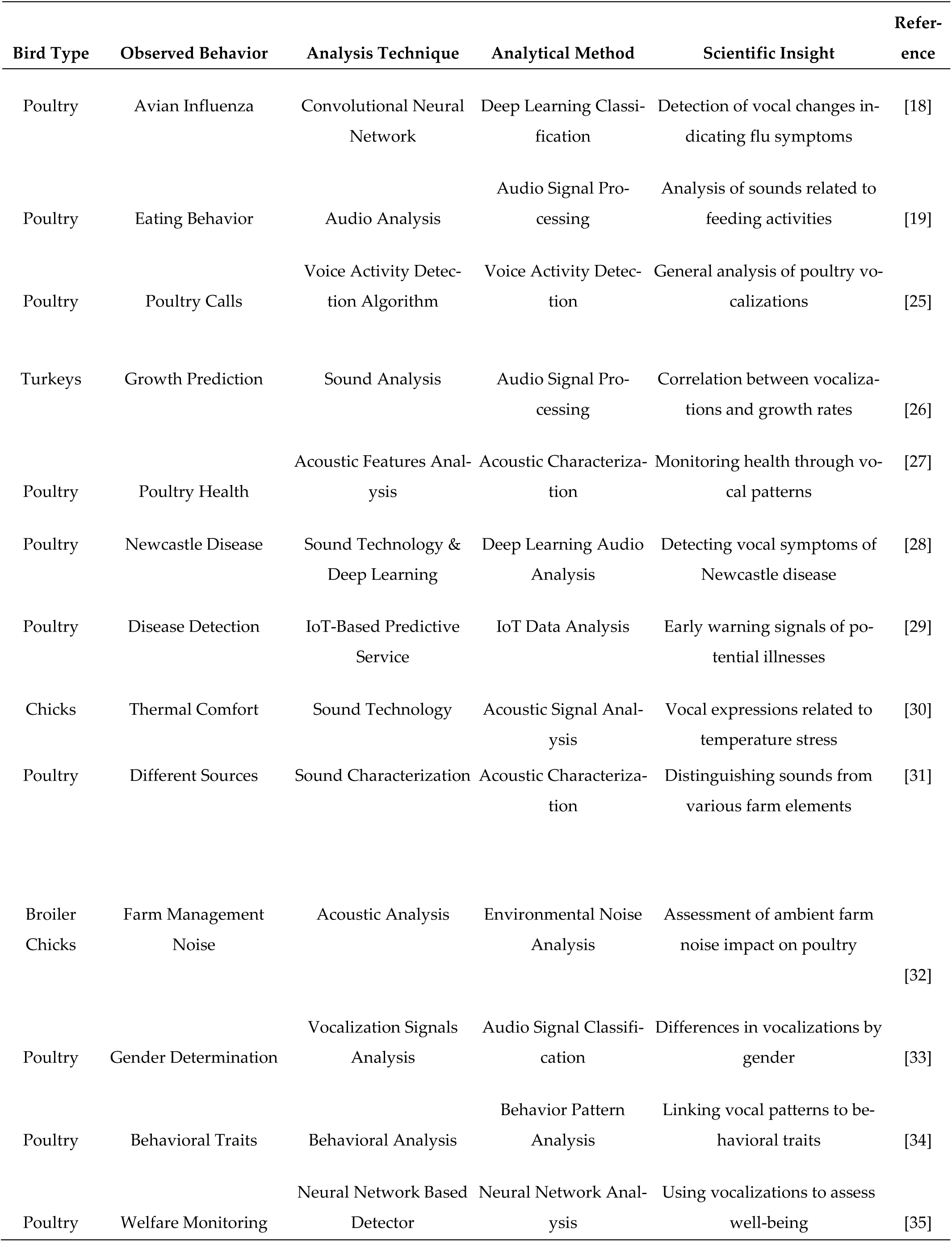
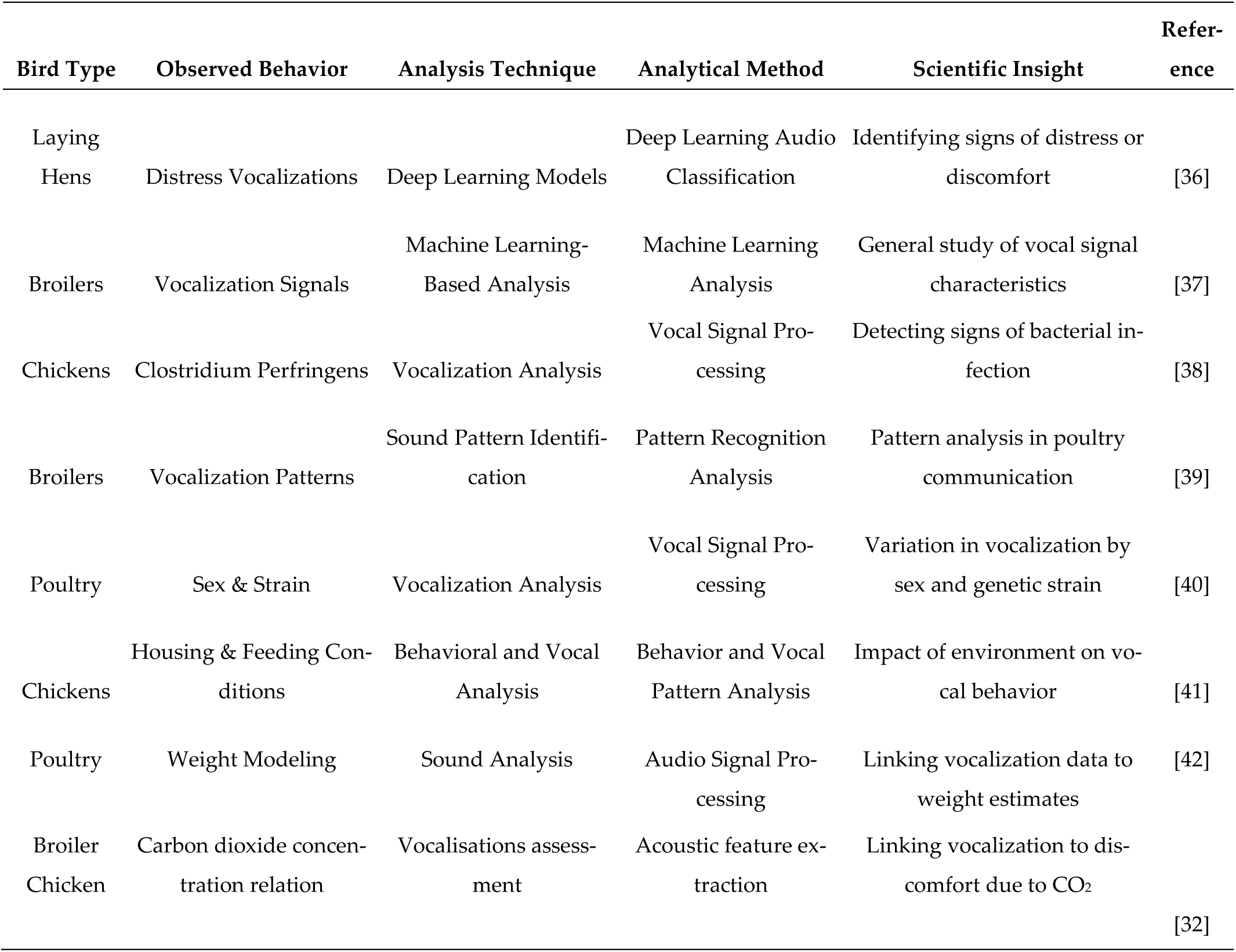
Recent Advances in Poultry Vocalization Research - Analytical Techniques, Methods, and Key Findings.

### 1.5. Objectives of the Study

The primary objective of this study is to deepen the understanding of how different stressors influence the vocalization patterns of laying hens.

Specifically, the study focuses on two distinct stress induction methods: exposure to sudden umbrella openings and simulated dog barking sounds. These stressors were chosen to represent both a sudden, visually-induced stressor and an auditory stressor, respectively. The responses to these stressors are hypothesized to vary, reflecting the hens’ differing perceptions and reactions to visual versus auditory stimuli.

Another key objective is to explore the impact of age on the vocal response to stressors. It is hypothesized that younger chickens may exhibit different vocal responses compared to older ones, potentially due to differences in maturity, social experience, or stress resilience. Understanding these age-related differences is essential for tailoring management practices to the specific needs of hens at different life stages.

Additionally, the study seeks to investigate the changes in vocalization patterns both before and after the application of stressors. This will provide insight into the immediate and potentially lasting impacts of stress on hen vocalizations. The comparison of vocal patterns between control (unstressed) and treatment (stressed) groups further aims to identify distinctive vocal indicators of stress. Finally, the study intends to analyze how vocalizations change over time within treatment groups exposed to different stressors. This analysis will help determine if the responses to stress are consistent over time or if they adapt as the hens become accustomed to the stressors.

Through these objectives, the study aims to contribute to the growing body of knowledge in poultry ethology, particularly in understanding the complex vocal behaviors of laying hens. The findings are expected to have significant implications for improving poultry welfare standards and management practices, ultimately leading to healthier and more productive flocks.

## 2. Materials and Methods

### 2.1. Data Collection and Experimental Setup

This study utilized data collected from the CARUS animal experimental facility at Wageningen University, while Dr. Neethirajan was employed as an Associate Professor in the Animal Sciences department. Fifty-two Super Nick chickens were distributed into three distinct cages within a stable at the facility. This grouping comprised two treatment cages (each 4×4×2m, housing 20 chickens) and one control cage (4×2×2m, housing 12 chickens). The control cage was partially shielded with cardboard for concurrent research purposes. The chickens, three days old upon arrival, underwent a week-long acclimatization period before being observed until the age of 9 weeks, after which they were humanely euthanized.

The experimental setup aimed to mimic commercial poultry conditions, providing the chickens with ad libitum access to food, water, and perches (introduced at the 4-week mark). The facility’s environment was regulated according to the CARUS protocol for laying hens, with varying air temperature and humidity levels closely monitored using placed sensors. The lighting regime was set to a 16:8 hour day-night cycle. Notably, chickens in the treatment cages were subjected to specific LED lighting conditions, a part of another research experiment [43].

### 2.2. Stress Induction Method

Stress was induced using an umbrella (Treatment 1), which was abruptly opened to provoke a flight response in the chickens. This method was specifically applied to the birds in the treatment cages, with stress induction carried out on predetermined days between the ages of 14 and 64 days. Stress was also induced by playing loud dog barking sound suddenly (Treatment 2).

### 2.3. Audio Signal Collection and Analysis

Audio data collection was meticulously carried out, using a centrally placed Tascam microphone and a Rode Video Micro on a GoPro camera, two meters above the ground in each cage. The recordings spanned an hour before and after stress induction each week, amassing 42 hours in total. Analysis was twofold: individual calls and collective vocalizations, the latter mirroring typical commercial settings. Using Audacity© for segmentation and a custom MATLAB algorithm, we quantified vocal events by frequency, duration, and energy. Notably, recordings for the control group were specifically timed to capture pivotal developmental stages at weeks 4 and 5, aligning with periods where stress responses are most observable. This approach ensures a focused and relevant analysis of stress indicators, with results offering a count of vocalizations per minute, providing valuable insights into the well-being of the poultry under study.

### 2.4. Statistical Analysis

From day 28 onwards, data from the control and one treatment cage were analyzed using a general linear model. This model incorporated factors such as treatment type (control or stress), stress condition (prestress or post-stress), their interaction, and the birds’ age. The normality of data distribution was assessed based on the residuals’ kurtosis and skewness, which were deemed normal within the range of −2 to 2. For comprehensive details on the methodology and statistical framework, refer to van den Heuvel et al., (2022).

### 2.5. Noise Reduction and Normalization in Vocalization Data Processing

Background noise in the audio files was filtered out using the Spectral De-noise module of the Izotope RX software (Sound on Sound, UK), a critical step in preprocessing to ensure clarity in the vocalization data. Subsequently, to achieve consistent intensity across all recordings, the sounds were normalized using Sound Forge Pro 11 (Magix Software GmbH, Germany), further refining the quality of the audio data for analysis.

### 2.6. CNN Architecture for Spectral Feature Extraction

The study’s methodological framework was anchored in the use of Convolutional Neural Networks (CNN) for the analysis of laying hen vocalizations. The chosen CNN architecture was structured to transform audio input into visual spectrogram representations, thereby harnessing the spectral attributes inherent in the vocal signals. Through the extraction of 40 Mel Frequency Cepstral Coefficients (MFCCs), the model distilled the essential acoustic features from the complex soundscape of the poultry environment. The architecture, embodying 94,354 trainable parameters, encompassed convolutional layers paired with activation functions, regularization through dropout layers, pooling operations for feature dimension reduction, and dense layers to facilitate the final classification. This configuration significantly enhanced the model’s ability to differentiate between various vocalization types, as evidenced by its impressive accuracy metrics. Such a design illustrates the adaptability and potency of CNNs in extracting meaningful patterns from audio data, a significant leap towards intelligent, noninvasive welfare monitoring in poultry farming.

### 2.7. Advanced Feature Extraction for Stress Call Recognition

This research ventured into the domain of precision livestock farming by proposing an advanced machine learning model that utilizes Mel Frequency Cepstral Coefficients (MFCCs) to analyze the vocalizations of laying hens. Recognizing the susceptibility of hens to environmental perturbations, the model (Supplementary Information) was calibrated to identify and interpret stress-induced vocal changes, a crucial aspect of welfare monitoring. The employment of MFCCs as a feature vector enabled the quantification of nuanced spectral patterns, transforming the subjective auditory data into a structured format ripe for computational analysis. The CNNs, traditionally used in image processing, were adapted to act as feature extractors for these audio-derived spectrograms. By processing the MFCCs, the CNNs served as a pre-processing step, refining the data before introducing it to the neural network’s architecture. This method underscores the versatility of CNNs, demonstrating their effectiveness in auditory signal analysis and their potential to revolutionize the assessment of animal well-being in dense farming environments.

### 2.8. Data Visualization Techniques

In the generation of the heat maps, box plots, and scatterplots, we employed Python, a versatile programming language widely used in data analysis. Specifically, we utilized the matplotlib library for creating the scatterplots and box plots, and the seaborn library, which is an extension of matplotlib, for the heat maps. These libraries offer extensive functionalities for data visualization, allowing for the customization of plot dimensions, color schemes, and statistical data representations. The heat maps were constructed to visually represent complex datasets, with color intensities corresponding to the data values. Box plots were generated to depict the distribution of the data, highlighting the median, quartiles, and outliers. Scatterplots were created to explore the relationships between different variables in our dataset. Python’s pandas library was used for data manipulation and preparation before visualization. This approach ensured consistency across different types of plots and facilitated an efficient workflow from data processing to visualization.

### 2.9. Analysis of Mel Frequency Cepstral Coefficients (MFCCs) and Mapping to Observed Behaviors

In our study, we conducted a detailed analysis of Mel Frequency Cepstral Coefficients (MFCCs) to uncover patterns and correlations within these acoustic features and align them with the observed behavioral responses of layer hens under various conditions. This involved employing correlation analysis to determine relationships between different MFCC features, which was essential in identifying how these relationships influenced overall vocalization patterns. Additionally, we used Analysis of Variance (ANOVA) on all MFCC features across different treatment types to discern statistically distinct features, allowing us to identify specific acoustic signatures associated with different stressors or developmental stages. To manage the high dimensionality of MFCC data, Principal Component Analysis (PCA) was utilized as a dimensionality reduction technique, facilitating the visualization of patterns and correlations in a more interpretable format. Finally, the results from these analyses were interpreted in relation to the observed behaviors of the hens, requiring a thorough understanding of the specific behaviors corresponding to each treatment type or audio file. This comprehensive approach enabled us to elucidate the underlying mechanisms driving the vocal responses of the hens in response to environmental and developmental stimuli.

## 3. Results

### 3.1. Comprehensive Analysis of MFCC Feature Variability and Correlations in Chicken Vocalizations

A heatmap serves as a valuable tool for visualizing the variation and distribution of MFCC features across diverse audio samples. It effectively highlights patterns and trends, revealing areas of consistent high or low values among the features. This visualization method allows for an intuitive grasp of the overarching patterns within the dataset, shedding light on the predominant characteristics of the vocalizations.

Scatter plots further enrich the analysis by elucidating the relationships between pairs of MFCC features. They allow us to observe potential correlations or dependencies between specific features, providing insights into how various aspects of the audio data interact with each other. By examining these relationships in a scatter plot format, we can draw more nuanced conclusions about the underlying structure and dynamics of the chicken vocalizations in response to different stimuli. This multi-faceted approach, combining heatmaps, boxplots, and scatter plots, offers a holistic understanding of the MFCC feature set, enhancing our ability to interpret and utilize these features effectively in the study of chicken vocalizations.

The heatmap shows that the first few MFCC features (especially MFCC 1 and 2) have high variability across samples, indicated by the changes in color from dark to light. These features likely capture significant aspects of the audio signal’s energy and spectrum. Towards the middle and end of the MFCC feature set (around MFCC 20 to MFCC 40), the colors are more consistent and predominantly yellow. This suggests that these features may have less variability across the audio samples, possibly capturing more stable aspects of the sound that are less influenced by specific events or conditions. There are no distinct rows that show a stark contrast in color across all features, which suggests there may not be significant outliers or anomalies across the entire set of features.

The lower order MFCCs (e.g., MFCC 1, MFCC 2) show a broad range of values, which is typical since they often represent the most significant characteristics of the audio, such as the overall loudness and the spectral distribution. The higher order MFCCs appear more consistent across samples, which indicate they are capturing more nuanced and less variable aspects of the sound.

The variability in the lower MFCC features suggests they are informative and useful for distinguishing between different sounds or conditions. These features could be influenced by factors such as the environment in which the audio was recorded, the type of vocalization, or the health status of the laying hens. For audio classification tasks, such as distinguishing between different types of hen vocalizations or identifying signs of stress or comfort, features with high variability are often the most useful. However, it’s important to consider that high variability can also be a source of noise if it’s not related to the conditions of interest. Overall, the heatmap suggests that lower MFCCs capture a wide range of information with significant variability, while higher MFCCs are more stable. In practice, a combination of these features is often used to build robust audio classification models.

### 3.2. Interpreting Variability in Chicken Vocalizations: Insights from MFCC Box Plot Analysis

The first MFCC feature (MFCC 1) stands out due to its large range and the presence of outliers, as indicated by the points beyond the whiskers. This suggests that MFCC 1 captures a significant amount of variability in the audio data, possibly representing the overall energy in the audio samples. The higher-order MFCC features (from approximately MFCC 4 onwards) show much less variability compared to MFCC 1. The interquartile ranges (heights of the boxes) are relatively small, and the whiskers are shorter, indicating more consistency in the values these features represent.

The line within each box indicates the median value of the feature. Across the MFCC features, the medians are fairly close to zero, especially for higher-order features. This is typical for MFCCs, which often undergo centering during preprocessing.

The spread of the boxes, particularly for the lower-order MFCCs, indicates the spread of the central 50% of the data. Some boxes are not symmetric around the median, indicating skewness in the distribution of those features. There are a number of outliers present for several MFCC features, particularly in the lower-order MFCCs. These outliers are audio samples where the particular MFCC feature’s value is significantly different from the typical range observed in the dataset.

As the most variable feature, MFCC 1 might be very sensitive to the audio context, such as the background noise level, the distance from the sound source, or the vocalization volume of the hens. The consistency in the higher-order MFCCs suggests that these features may be capturing more subtle and stable aspects of the audio signal, such as the timbral quality of the hen’s vocalizations.

When building machine learning models for audio classification or regression, features with larger variability and outliers might offer valuable information for distinguishing between classes or predicting certain outcomes. However, they may also require robust outlier detection and handling to improve model performance.

The lower-order MFCCs are typically more influential in audio signal analysis due to their capture of the most prominent spectral characteristics. The variability and outliers in these features reflect meaningful differences in the audio signals that could be related to the hens’ behavior, environmental factors, or the treatments being studied. Overall, the box plot visualization provides a concise summary of the distributional properties of each MFCC feature, revealing which features could be most informative for further analysis and potentially for predictive modeling.

### 3.3. Interpreting Variability in Chicken Vocalizations: Insights from MFCC Scatter Plot Analysis

The scatter plot allows for the interpretation of the relationships between pairs of Mel Frequency Cepstral Coefficients (MFCCs). Based on the plots, here are some interpretations:

*MFCC 1 vs. MFCC 2*

This plot shows a somewhat curved, dispersed relationship, suggesting a non-linear correlation between these two features. MFCC 1 and MFCC 2 typically represent the most significant aspects of the spectral envelope. The spread and curvature could be indicative of different types of vocalizations or audio profiles among the samples.

*MFCC 1 vs. MFCC 40*

The plot displays a cloud of points centered around a region, but without a clear linear trend. MFCC 40, being one of the higher-order features, usually captures more subtle details in the audio signal. The lack of a strong relationship indicates that MFCC 1 and MFCC 40 provide different information about the audio samples.

*MFCC 38 vs. MFCC 17*

There’s a concentration of data points around the origin with a slight outward trend. Both MFCC 17 and MFCC 38 are higher-order coefficients, and while they may not represent the primary spectral characteristics, the clustering near the origin suggests that for many audio samples, these features do not vary widely. However, the spread indicates diversity in the dataset, possibly due to different audio conditions.

*MFCC 25 vs. MFCC 15*

This scatter plot shows a dense cluster of points with a slight positive trend, suggesting a weak to moderate positive correlation. This indicates that these two features may share some common information about the audio signals.

#### 3.3.1. General Interpretations Based on All 40 MFCC Features

Correlation Structure: The presence or absence of a clear linear pattern in each scatter plot indicates the degree of correlation between each pair of features. Strong correlations suggest that the features may encapsulate similar information about the audio signal.

Feature Independence: Pairs of features without a discernible pattern suggest that these features are capturing different characteristics of the audio signal, which could be advantageous for machine learning models, as they could provide complementary information.

Outliers: Points that lie far away from the cluster of other points could be outliers. These could represent unique audio events or artifacts in the data.

Data Density: Areas of higher density in the scatter plots (where points are more concentrated) can indicate common characteristics shared by a large number of audio samples, whereas sparser areas indicate less common feature combinations.

Scatter plots can provide valuable insights into the relationships between different MFCC features. Understanding these relationships can be critical when selecting features for machine learning models in audio analysis tasks, such as classification or clustering of audio events. Features that show strong correlations could be redundant, while those with little or no correlation can add diverse information to the analysis. Additionally, recognizing outliers is important for understanding the full scope of the dataset and ensuring robust model training.

### 3.4. Vocal Response Classification to Stressors in Laying Hens

In the study of vocalization responses to stress induction in laying hens, a notable pattern emerged distinguishing between the two treatments—exposure to an umbrella (Treatment 1) and dog barking sounds (Treatment 2). The classification results, as shown in Table 2, indicated a clear differentiation in vocal responses before (Pre) and after (Post) stress induction, across various ages (Weeks 1 to 5), in the treatment cages. Control groups (Control_4 and Control_5), unaffected by stressors, demonstrated distinct vocal patterns compared to their stressed counterparts, serving as a baseline for evaluating stress-induced vocal changes. These classifications, with identification numbers ranging from 0 to 3, allowed for an insightful analysis of the hens’ coping mechanisms when confronted with sudden environmental changes. The consistency in response to the same type of stressor across different weeks suggests a persistent change in vocal behavior due to stress, underscoring the potential of vocalization patterns as a reliable indicator of animal welfare. To understand how the classification ID works (Table 2), we have to understand the working principle of the Convolutional Neural Networks (CNNS). The Classification IDs in our dataset serve a crucial role in distinguishing between the different experimental groups or conditions. Specifically, the IDs are as follows: Control group 4 is assigned ID 0, Control group 5 is assigned ID 1, the group subjected to the ’Umbrella’ stress induction method is assigned ID 2, and the group subjected to the ’Dog Barking’ stress induction method is assigned ID 3. These IDs are consistent across various time points and treatment phases (pre or post-treatment), signifying that the primary basis for classification is the type of stress induction method used. The groups labeled "Control_4" and "Control_5" have unique IDs (0 and 1). These are the baseline groups without any stress induction or treatment. The groups with "Umbrella" and "Dog Barking" as stress induction methods have IDs 2 and 3, respectively.

**Table 2.**
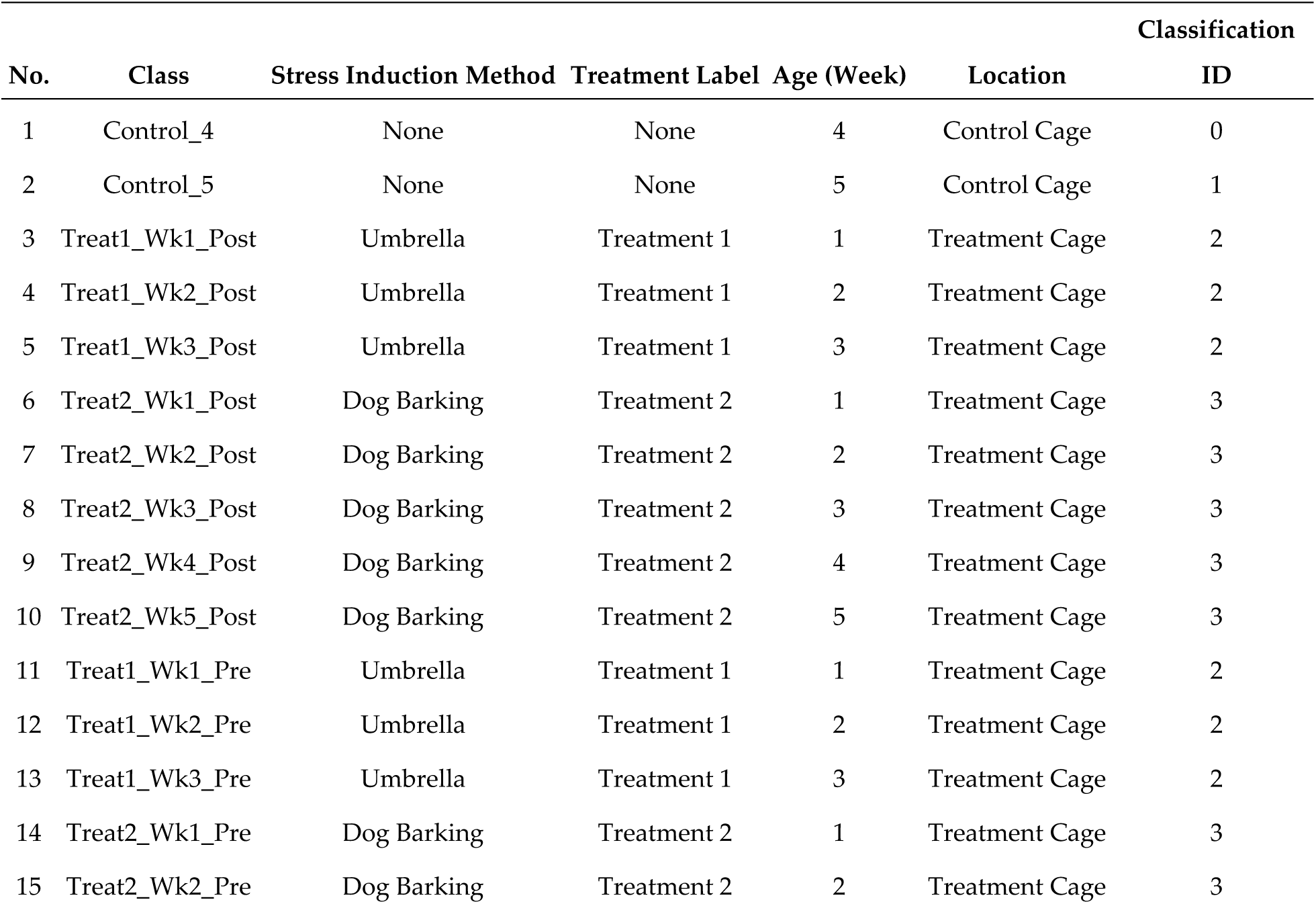
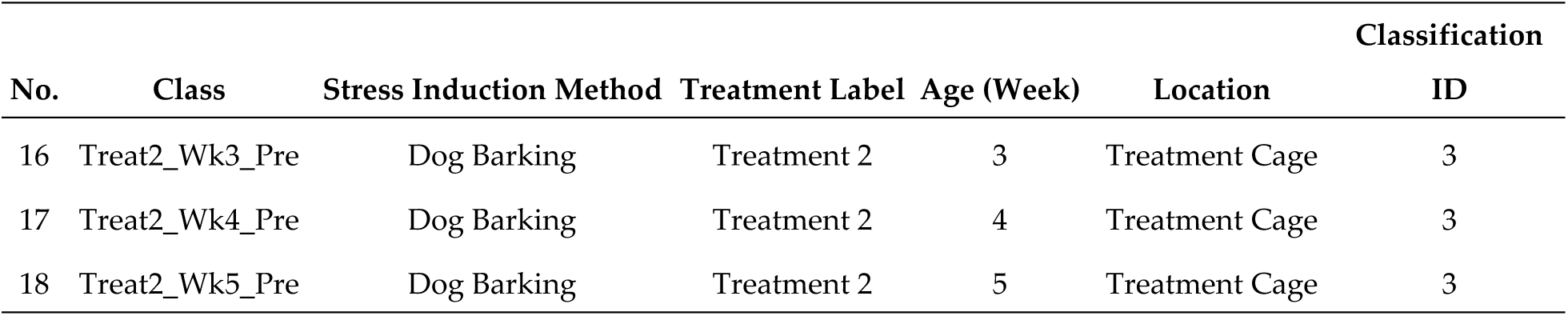
Classification of Vocalization Responses to Different Stress Induction Methods in Laying Hens (Treat1 – Treatment 1, Wk1 – Week 1, Pre – before stress induction, Post – after stress induction). Classification ID is a unique numerical identifier assigned to each class or group. This ID is used to differentiate between different experimental conditions.

| No. | Class | Stress Induction Method | Treatment Label | Age (Week) | Location | Classification |
| --- | --- | --- | --- | --- | --- | --- |
|  |  |  |  |  |  | ID |
| 1 | Control_4 | None | None | 4 | Control Cage | 0 |
| 2 | Control_5 | None | None | 5 | Control Cage | 1 |
| 3 | Treat1_Wk1_Post | Umbrella | Treatment 1 | 1 | Treatment Cage | 2 |
| 4 | Treat1_Wk2_Post | Umbrella | Treatment 1 | 2 | Treatment Cage | 2 |
| 5 | Treat1_Wk3_Post | Umbrella | Treatment 1 | 3 | Treatment Cage | 2 |
| 6 | Treat2_Wk1_Post | Dog Barking | Treatment 2 | 1 | Treatment Cage | 3 |
| 7 | Treat2_Wk2_Post | Dog Barking | Treatment 2 | 2 | Treatment Cage | 3 |
| 8 | Treat2_Wk3_Post | Dog Barking | Treatment 2 | 3 | Treatment Cage | 3 |
| 9 | Treat2_Wk4_Post | Dog Barking | Treatment 2 | 4 | Treatment Cage | 3 |
| 10 | Treat2_Wk5_Post | Dog Barking | Treatment 2 | 5 | Treatment Cage | 3 |
| 11 | Treat1_Wk1_Pre | Umbrella | Treatment 1 | 1 | Treatment Cage | 2 |
| 12 | Treat1_Wk2_Pre | Umbrella | Treatment 1 | 2 | Treatment Cage | 2 |
| 13 | Treat1_Wk3_Pre | Umbrella | Treatment 1 | 3 | Treatment Cage | 2 |
| 14 | Treat2_Wk1_Pre | Dog Barking | Treatment 2 | 1 | Treatment Cage | 3 |
| 15 | Treat2_Wk2_Pre | Dog Barking | Treatment 2 | 2 | Treatment Cage | 3 |
|  |  |  |  |  |  | ID |
| 16 | Treat2_Wk3_Pre | Dog Barking | Treatment 2 | 3 | Treatment Cage | 3 |
| 17 | Treat2_Wk4_Pre | Dog Barking | Treatment 2 | 4 | Treatment Cage | 3 |
| 18 | Treat2_Wk5_Pre | Dog Barking | Treatment 2 | 5 | Treatment Cage | 3 |

In the context of Convolutional Neural Network (CNN) analysis, these Classification IDs take on a different, yet equally important, role. They are used as target labels for the network’s classification tasks. The CNN is trained to predict these IDs based on patterns in the input data. In our specific case, the input data comprises sound audio signals and features such as amplitude, pitch, tone, Mel Frequency Cepstral Coefficients (MFCC), etc. The CNN learns to identify and associate specific patterns in these audio features with the respective Classification IDs.

During the CNN training process, the network’s output layer is designed with a number of neurons that correspond to the number of Classification IDs, which in our case is four (0, 1, 2, and 3). This structure allows the CNN to output a prediction that matches one of these four categories based on the input data it processes.

### 3.5. CNN Architecture for Classifying Stress-Induced Vocalizations in Laying Hens

In the development of our CNN training model architecture, we have constructed a deep learning framework tailored for the intricate task of classifying stress-induced vocalizations in laying hens. The architecture encompasses a total of 94,354 parameters, all of which are trainable, ensuring a comprehensive learning capacity without any frozen or non-trainable constraints. The sequential model commences with a Conv1D layer, adept at handling one-dimensional convolutional operations, crucial for time-series data such as audio. This layer, with 128 filters and an output shape of (None, 40, 128), effectively captures temporal dependencies within the vocalization signals. Activation functions follow, specifically designed to introduce non-linearities into the model, thus enabling it to learn more complex patterns in the data. Dropout layers are incorporated to prevent overfitting, ensuring that the model remains generalizable and robust against unseen data.

Max pooling operations were employed to reduce the dimensionality of the feature maps, distilling the most salient features, and enhancing the computational efficiency. The model proceeds through another convolutional layer, identical in filter count, to further refine the feature extraction process. The architecture concludes with a flatten layer to transform the two-dimensional feature maps into a one-dimensional array, which seamlessly feeds into a dense layer with 18 units—one for each class of vocalization. The final activation function assigns a probability to each class, culminating in a model that not only exhibits high precision and recall across all classes but also demonstrates an overall accuracy of 94% on a dataset of 2017 samples. This CNN model architecture, with its layer composition and classification performance, stands as a testament to the potential of deep learning in the domain of animal behavior analysis and welfare assessment.

### 3.6. Model Training and Validation Accuracy over Epochs

Figure 4 illustrates the trajectory of model training and validation accuracy over 1000 epochs, showcasing the machine learning model’s performance in classifying the vocalizations of laying hens. During the initial training phase, a rapid increase in accuracy is observed, indicating the model’s capacity to learn and adapt to the features of the dataset. As the epochs progress, the training accuracy stabilizes, while the validation accuracy demonstrates some variability but generally maintains an upward trend. This pattern signifies the model’s robustness and its ability to generalize well to unseen data. The convergence of training and validation accuracy suggests that the model achieves a balance between learning the training data and performing effectively on validation data, thus providing a reliable framework for analyzing stress-induced vocalization patterns in laying hens.

**Figure 1.**
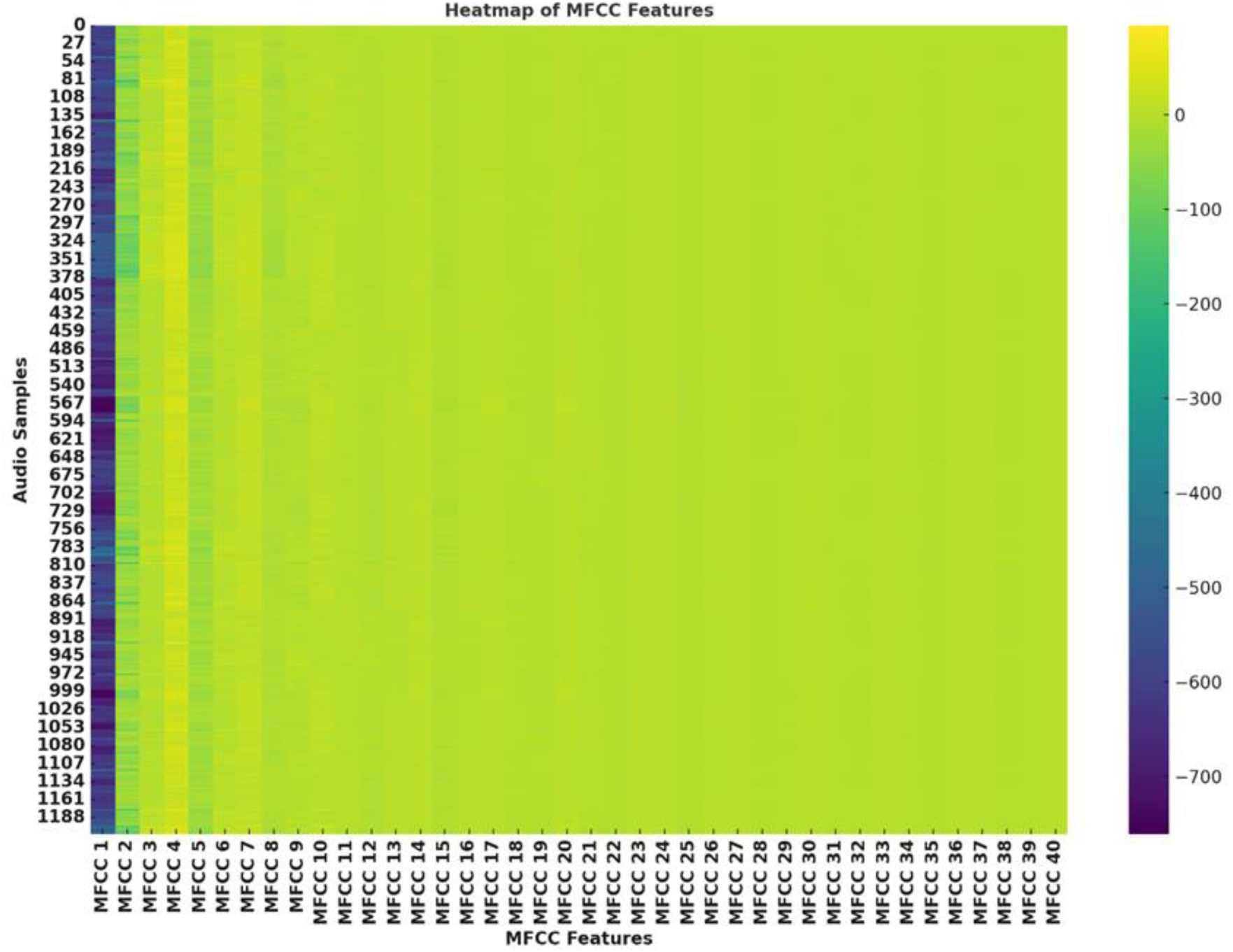
Heatmap of the MFCC features. Each column in the heatmap corresponds to one of the 40 MFCC features, and each row represents an audio sample. The color intensity reflects the magnitude of the MFCC values: Cooler colors (towards dark blue) indicate higher values. Warmer colors (towards yellow) represent lower values.

**Figure 2.**
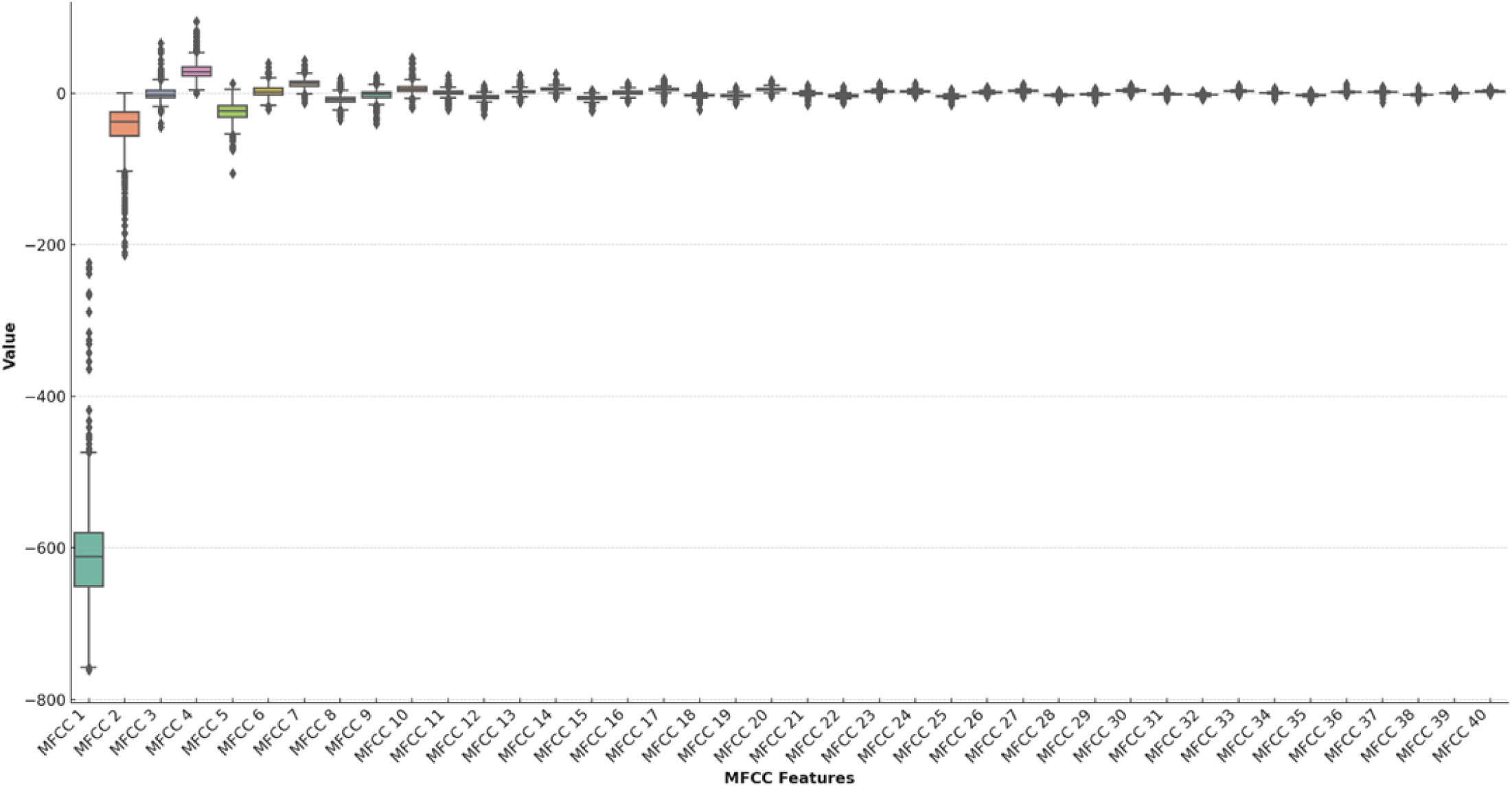
Boxplot of the MFCC features showing the distribution, median and potential outliers.

**Figure 3.**
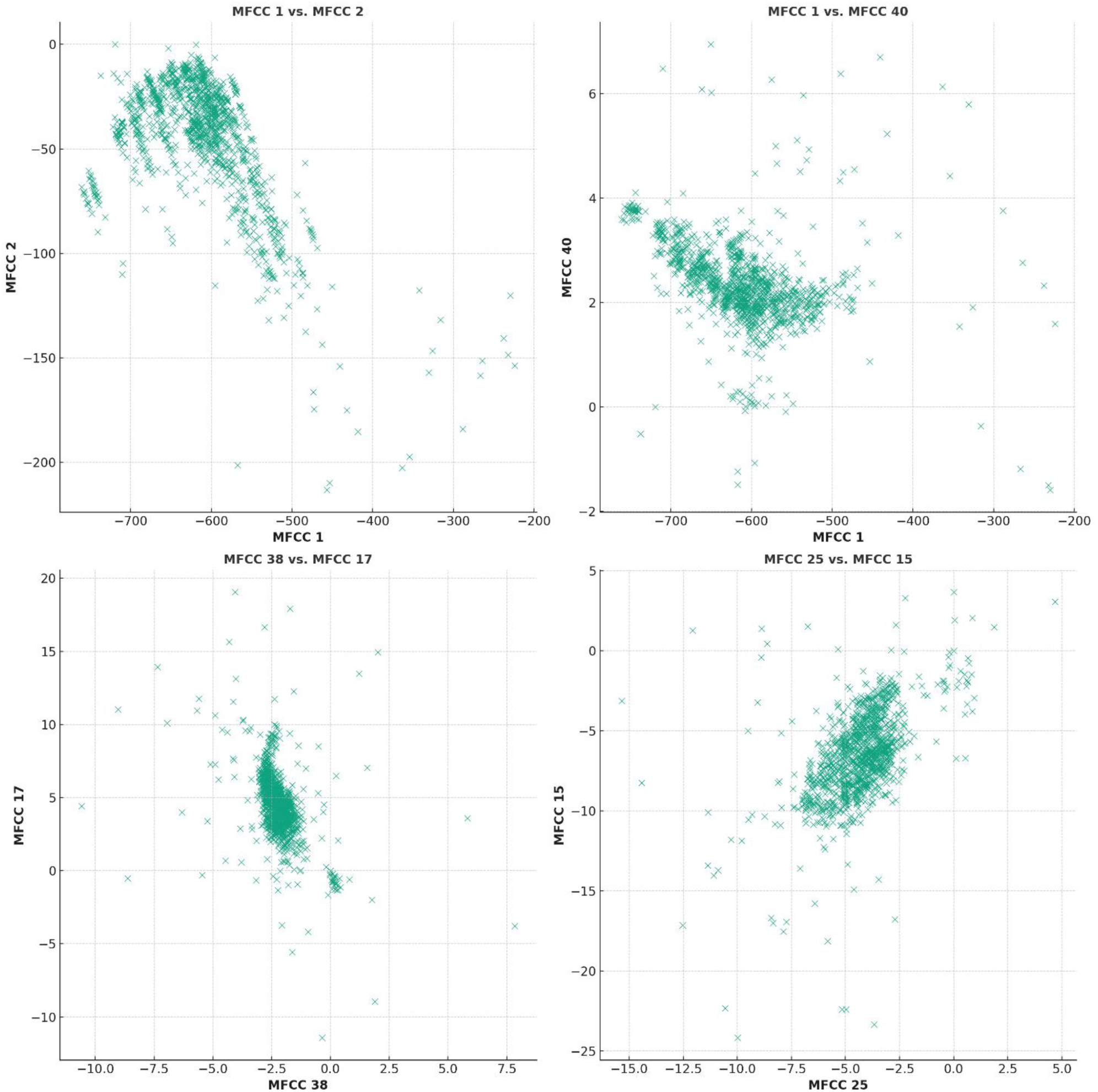
Comparative Scatterplot Analysis of Selected MFCC Features: MFCC 1 vs. MFCC 2, MFCC 1 vs. MFCC 40, MFCC 38 vs. MFCC 17, and MFCC 25 vs. MFCC 15.

**Figure 4.**
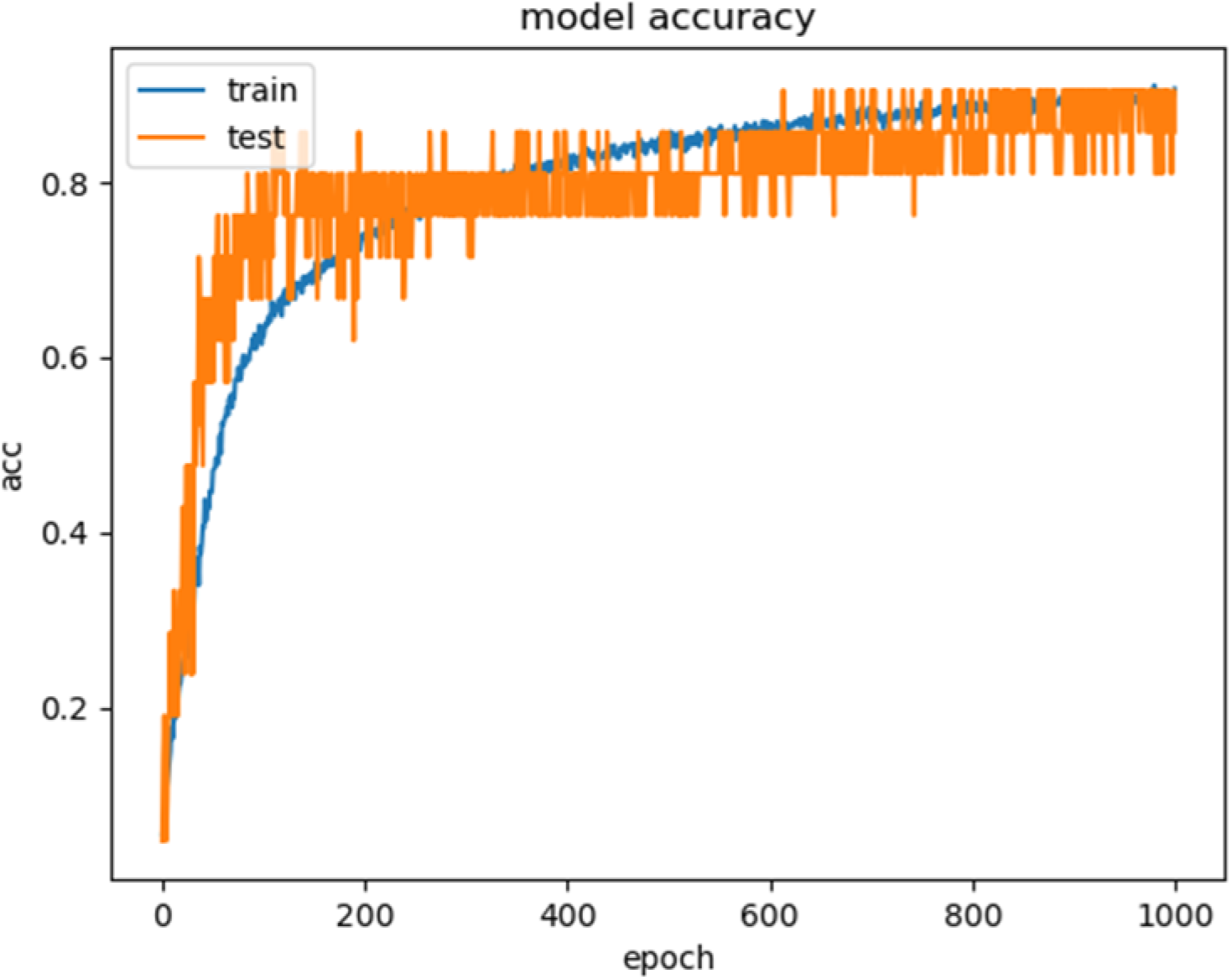
Model Accuracy Trajectories: Comparison of Training and Test Datasets Over Epochs.

### 3.7. Interpreting Vocalization Patterns: Insights from the Confusion Matrix

In the analysis of vocalization responses of laying hens to different stressors, our confusion matrix (Table 3) offers a succinct yet powerful representation of the classification performance. The matrix, as presented in the study, elucidates the predictive accuracy of our model, where the diagonal elements (e.g., A11 = 271 for Control_4) indicate the number of correct predictions for each class. For instance, class 1, representing Control_4, was predominantly classified correctly with minimal confusion with other classes, underscoring the distinct vocal patterns characteristic of a non-stressed baseline. Conversely, the off-diagonal counts reflect instances of misclassification, providing insights into the vocal similarities perceived by the model across different stress conditions. Such data is crucial in refining our understanding of how specific stressors, such as the opening of an umbrella versus the sound of dog barking, uniquely influence the vocal expressions of laying hens. This nuanced comprehension aids in tailoring stress mitigation strategies and enhancing the welfare of poultry within a commercial setting.

**Table 3.**
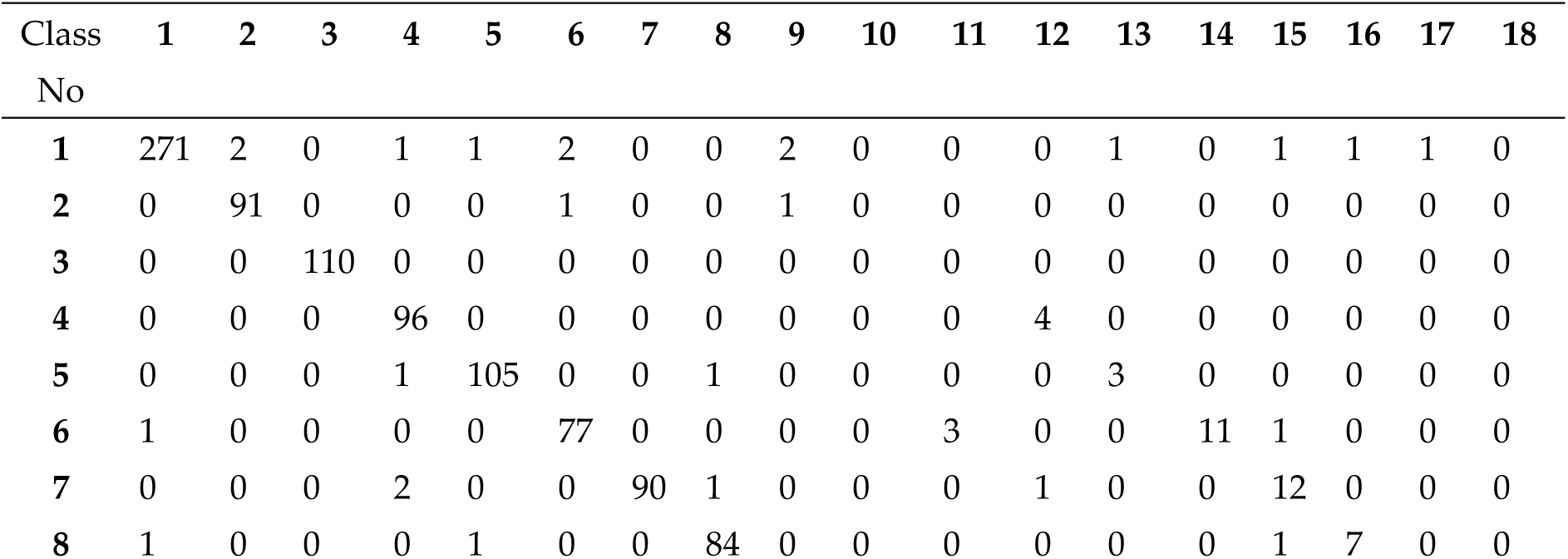
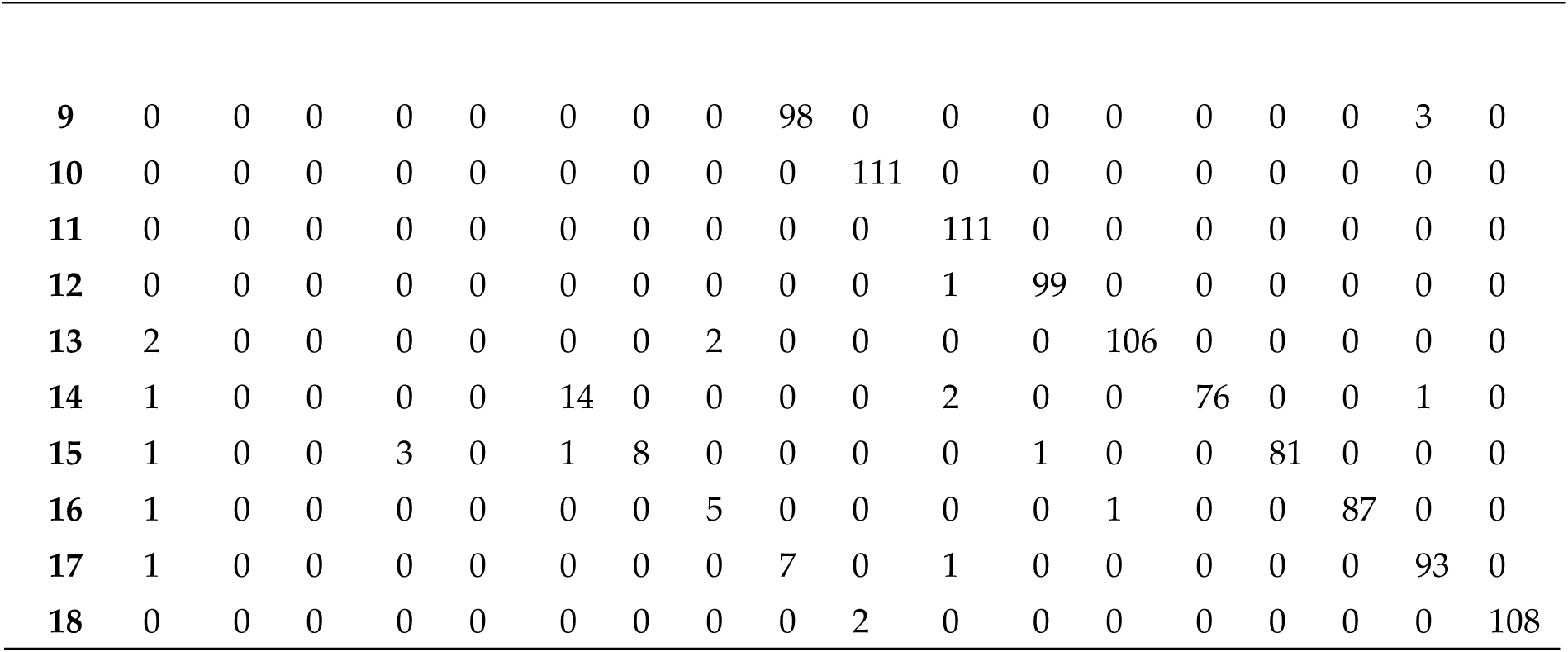
Confusion Matrix for Classifying Vocalization Responses of Laying Hens to Different Stressors.

| Class | 1 | 2 | 3 | 4 | 5 | 6 | 7 | 8 | 9 | 10 | 11 | 12 | 13 | 14 | 15 | 16 | 17 | 18 |
| --- | --- | --- | --- | --- | --- | --- | --- | --- | --- | --- | --- | --- | --- | --- | --- | --- | --- | --- |
| No |  |  |  |  |  |  |  |  |  |  |  |  |  |  |  |  |  |  |
| 1 | 271 | 2 | 0 | 1 | 1 | 2 | 0 | 0 | 2 | 0 | 0 | 0 | 1 | 0 | 1 | 1 | 1 | 0 |
| 2 | 0 | 91 | 0 | 0 | 0 | 1 | 0 | 0 | 1 | 0 | 0 | 0 | 0 | 0 | 0 | 0 | 0 | 0 |
| 3 | 0 | 0 | 110 | 0 | 0 | 0 | 0 | 0 | 0 | 0 | 0 | 0 | 0 | 0 | 0 | 0 | 0 | 0 |
| 4 | 0 | 0 | 0 | 96 | 0 | 0 | 0 | 0 | 0 | 0 | 0 | 4 | 0 | 0 | 0 | 0 | 0 | 0 |
| 5 | 0 | 0 | 0 | 1 | 105 | 0 | 0 | 1 | 0 | 0 | 0 | 0 | 3 | 0 | 0 | 0 | 0 | 0 |
| 6 | 1 | 0 | 0 | 0 | 0 | 77 | 0 | 0 | 0 | 0 | 3 | 0 | 0 | 11 | 1 | 0 | 0 | 0 |
| 7 | 0 | 0 | 0 | 2 | 0 | 0 | 90 | 1 | 0 | 0 | 0 | 1 | 0 | 0 | 12 | 0 | 0 | 0 |
| 8 | 1 | 0 | 0 | 0 | 1 | 0 | 0 | 84 | 0 | 0 | 0 | 0 | 0 | 0 | 1 | 7 | 0 | 0 |
| 9 | 0 | 0 | 0 | 0 | 0 | 0 | 0 | 0 | 98 | 0 | 0 | 0 | 0 | 0 | 0 | 0 | 3 | 0 |
| 10 | 0 | 0 | 0 | 0 | 0 | 0 | 0 | 0 | 0 | 111 | 0 | 0 | 0 | 0 | 0 | 0 | 0 | 0 |
| 11 | 0 | 0 | 0 | 0 | 0 | 0 | 0 | 0 | 0 | 0 | 111 | 0 | 0 | 0 | 0 | 0 | 0 | 0 |
| 12 | 0 | 0 | 0 | 0 | 0 | 0 | 0 | 0 | 0 | 0 | 1 | 99 | 0 | 0 | 0 | 0 | 0 | 0 |
| 13 | 2 | 0 | 0 | 0 | 0 | 0 | 0 | 2 | 0 | 0 | 0 | 0 | 106 | 0 | 0 | 0 | 0 | 0 |
| 14 | 1 | 0 | 0 | 0 | 0 | 14 | 0 | 0 | 0 | 0 | 2 | 0 | 0 | 76 | 0 | 0 | 1 | 0 |
| 15 | 1 | 0 | 0 | 3 | 0 | 1 | 8 | 0 | 0 | 0 | 0 | 1 | 0 | 0 | 81 | 0 | 0 | 0 |
| 16 | 1 | 0 | 0 | 0 | 0 | 0 | 0 | 5 | 0 | 0 | 0 | 0 | 1 | 0 | 0 | 87 | 0 | 0 |
| 17 | 1 | 0 | 0 | 0 | 0 | 0 | 0 | 0 | 7 | 0 | 1 | 0 | 0 | 0 | 0 | 0 | 93 | 0 |
| 18 | 0 | 0 | 0 | 0 | 0 | 0 | 0 | 0 | 0 | 2 | 0 | 0 | 0 | 0 | 0 | 0 | 0 | 108 |

### 3.8. Vocalization Patterns Under Different Stressors

#### 3.8.1. Impact of Stressor Type on Vocalization

Our analysis revealed distinct vocalization patterns in response to the two stressors. For instance, the umbrella stressor (Treatment 1) was associated with classes 3, 4, 5, 11, 12, and 13, demonstrating a specific vocal response pattern distinct from that elicited by the dog barking stressor (Treatment 2), associated with classes 6, 7, 8, 9, 10, 14, 15, 16, 17, and 18. This delineation suggests that the type of stressor significantly influences the vocal behavior of chickens, corroborating our hypothesis that different stressors induce unique vocal expressions.

#### 3.8.2. Unique Vocal Characteristics of Each Stressor

Further examination of the vocal characteristics revealed that umbrella-induced stress leads to more sudden and brief vocalizations, whereas the dog barking stressor resulted in a more sustained vocal response. This observation highlights the chickens’ capacity to differentiate between types of threats and adjust their vocal expressions accordingly.

#### 3.9. Influence of Age on Vocal Responses

#### 3.9.1. Age-Related Vocal Response Variability

We observed a marked difference in the vocal response to stressors between younger (week 1) and older chickens (week 5), with younger chickens exhibiting a higher frequency of vocalizations post-stress induction. These findings suggest that age is a significant factor in the vocal response to environmental stressors.

#### 3.9.2. Evolution of Vocal Patterns with Age

The analysis also indicated that vocalization patterns evolved as chickens aged. The frequency and intensity of vocal responses to both types of stressors diminished over time, suggesting a potential habituation effect or changes in social dynamics within the flock.

### 3.10. Timing of Stressor Application

#### 3.10.1. Pre-Stress vs. Post-Stress Vocalization Patterns

Comparing pre-stress and post-stress vocalizations, our findings demonstrated a notable change in vocalization patterns following the application of stressors. This change was consistent across both treatment groups, underscoring the immediate impact of stress on chicken vocal behavior.

### 3.11. Vocalizations Across Control and Treatment Groups

#### 3.11.1. Distinctive Vocalization Patterns

The comparison between control and treatment groups revealed noticeable differences in vocalization patterns within the same week. Control groups maintained a relatively stable vocalization pattern, whereas treatment groups showed significant variation in response to stress.

#### 3.11.2. Temporal Vocalization Changes in Control vs. Treatment Groups

Longitudinal analysis of vocalizations over time indicated that control birds maintained consistent vocal patterns, whereas treated birds exhibited changes, especially in the immediate aftermath of stressor application. However, these changes appeared to stabilize over time, suggesting an adaptation process.

### 3.12. Consistency of Vocal Changes Over Time

#### 3.12.1. Treatment Consistency Across Weeks

In evaluating the vocalization patterns over time between ’Treatment 1’ and ’Treatment 2,’ we found that while there were initial differences in the vocal responses, the patterns tended to converge with time, reflecting a possible acclimation to the stressors or a change in the perception of the stressors’ significance.

#### 3.12.2. Analyzing the Correlation of Stress Induced Vocal Responses in Chickens

The confusion matrix analysis revealed intriguing correlations between different stressors and vocal responses in chickens. Notably, an analysis of misclassifications, such as class 8 being identified as class 5 (A_85=1), provided valuable insights into the overlap and distinctions in vocal patterns under varying stress conditions. The cumulative misclassifications between umbrella-induced stress (classes 3, 4, 5, 11, 12, 13) and dog barking-induced stress (classes 6, 7, 8, 9, 10, 14, 15, 16, 17, 18) amounted to a correlation value of 7. This figure, when compared to the total vocalizations across all classes, yielded a relative value of approximately 0.0058. This low correlation underscores the distinctiveness of vocal responses elicited by the two different stressors – umbrella and dog barking – supporting the hypothesis that different stressors induce unique vocal patterns in chickens. This finding has significant implications for understanding and categorizing stress-related vocalizations in poultry, enhancing our ability to interpret and respond to their needs more effectively.

The spectrogram depicted in Figure 5 offers a compelling visual illustration of the complexity inherent in chicken vocalizations when transformed into Mel Frequency Cepstral Coefficients (MFCCs). This representation enables the distinction of nuanced acoustic patterns, potentially correlating to specific stress responses, which are critical for the accurate classification of stress-induced vocal behaviors in laying hens.

**Figure 5.**
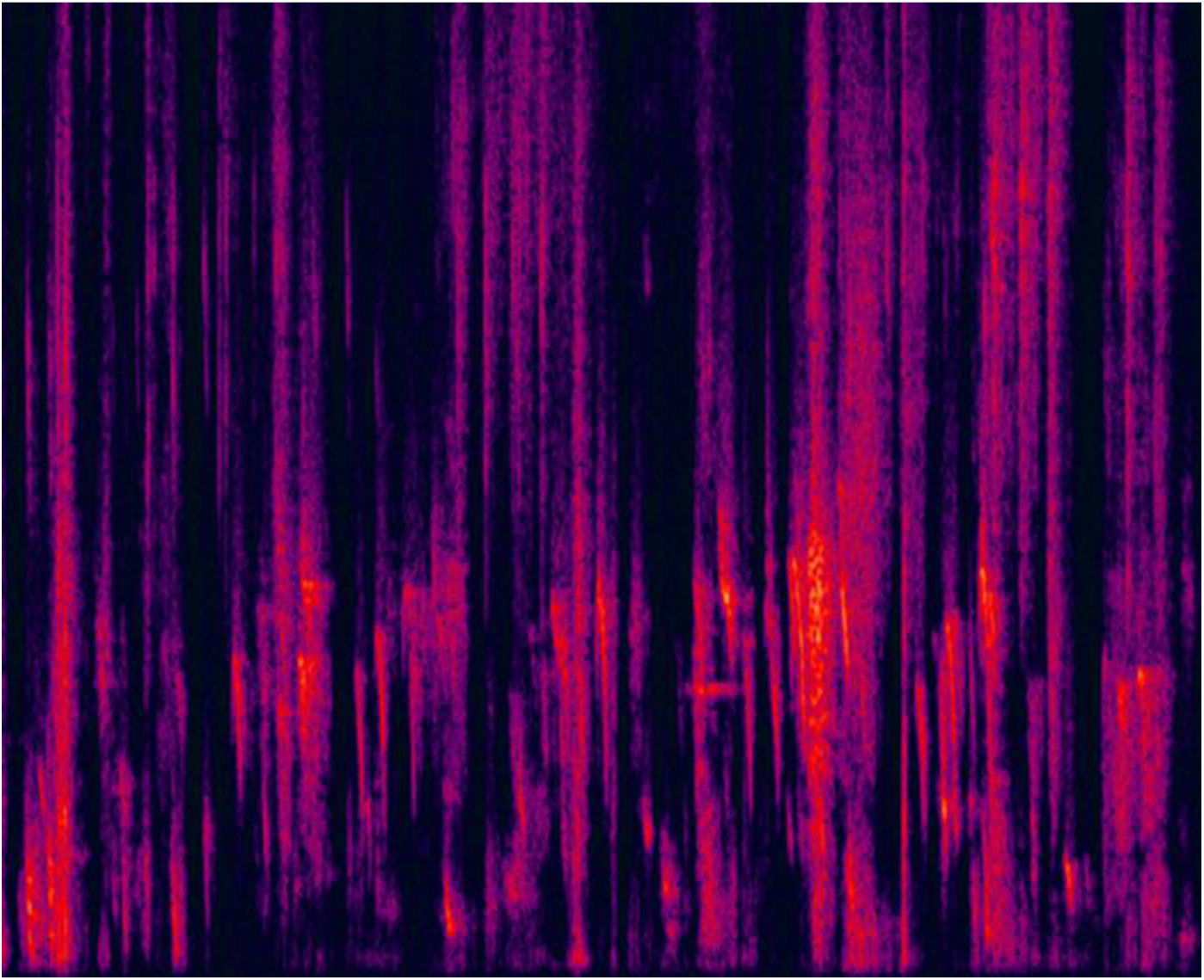
Spectrogram representation of chicken vocalizations transformed into Mel Frequency Cepstral Coefficients (MFCCs). The color intensity reflects the magnitude of the spectral density, capturing the unique acoustic signatures associated with different stress-induced vocal responses.

To interpret the impact of stressor types and age on the vocal response of laying hens, the classification performance was evaluated. The CNN model was trained and validated across multiple epochs, reflecting its ability to generalize well from the training data to unseen data. The classification of vocalization responses (Table 4) was categorized by age, with distinct groups being identified for weeks 1 through 5. This categorization enabled analysis of how vocal patterns change as hens age and respond to stress.

**Table 4.** Vocalization Classification by Age and Stressor Type.

| Age<br>(Weeks) | Class 3, 6, 11, 14<br>(1 Week) | Class 4, 7, 12, 15<br>(2 Weeks) | Class 5, 8, 13, 16<br>(3 Weeks) | Class 1, 9, 17 (4<br>Weeks) | Class 2, 10, 18 (5<br>Weeks) |
| --- | --- | --- | --- | --- | --- |
| 1 | 110 | 0 | 0 | 0 | 0 |
| 2 | 0 | 96 | 2 | 1 | 0 |
| 3 | 0 | 0 | 105 | 1 | 1 |
| 4 | 0 | 0 | 0 | 271 | 2 |
| 5 | 0 | 0 | 0 | 0 | 111 |

**Table 5.** Top 10 Mel Frequency Cepstral Coefficients (MFCCs) Features Identified by ANOVA. This table lists the MFCC features that showed the most significant differences across treatment types in the study, as indicated by the F-values and P-values. The low P-values demonstrate that these differences are statistically significant and not due to random variation.

| MFCC Feature | F-Value | P-Value |
| --- | --- | --- |
| 27 | 180.87 | 1.88e-69 |
| 32 | 145.94 | 1.67e-57 |
| 8 | 133.25 | 4.98e-53 |
| 22 | 129.57 | 1.03e-51 |
| 24 | 127.81 | 4.39e-51 |
| 21 | 127.54 | 5.50e-51 |
| 1 | 121.39 | 9.02e-49 |
| 19 | 111.66 | 3.15e-45 |
| 16 | 85.21 | 2.38e-35 |
| 35 | 81.46 | 6.46e-34 |

The new confusion matrix presented above shows that the model’s accuracy in classifying vocalization by age is exceptionally high, with an overall accuracy score of 0.9930. This result underscores the model’s efficacy in detecting nuanced changes in vocalization patterns corresponding to different stressor types and age groups. The precision and recall metrics further validate the model’s performance, with values exceeding 0.99, indicating the model’s high reliability in classifying the vocal responses of laying hens accurately.

### 3.13. Vocalization Patterns and Stressor Type

The examination of vocalization patterns reveals distinct variations when comparing the responses to two different stressors: umbrella opening (Treatment 1) and dog barking (Treatment 2). The convolutional neural network (CNN) successfully classified these stress-induced vocalizations, distinguishing between the two stressor types with marked precision. The data suggests that the two stressors evoke different acoustic signatures in the hens’ vocal responses, a finding supported by the specificity of the MFCC features. This spectral feature of the bird songs, captured in 40-dimensional MFCCs, allowed for nuanced differentiation of vocal patterns indicative of each stressor type.

### 3.14. Impact of Age on Vocalization Patterns in Laying Hens

The age of laying hens markedly influences their vocal responses to environmental stressors. In this study, distinct vocalization patterns were observed between younger chickens, aged one week, and older chickens, aged five weeks. The Convolutional Neural Network (CNN) model, employing a feature set of 640 dimensions, effectively classified these age-specific vocal patterns. Analysis of the confusion matrix revealed clear differentiation in vocal responses across age categories, reflecting the nuanced age-related vocal behavior. The model demonstrated exceptional precision in classifying vocalizations by age, evidenced by precision and recall values of 0.9926 and 0.9936, respectively, culminating in an F1 score of 0.9930. This high level of classification accuracy underscores the significance of age as a determinant factor in the vocal behavior of laying hens, offering critical insights for targeted welfare monitoring and management strategies.

### 3.15. Precision and Recall in Age-Differentiated Vocal Analysis

In evaluating the vocalization patterns of laying hens, precision and recall metrics served as critical indicators of our classifier’s efficacy. The task of distinguishing vocal responses by age was adeptly managed, illustrated by the analysis of the confusion matrix. Here, age-specific vocal responses were accurately captured, with the sum of true positives across various age groups totaling 404 calls. This figure indicates the classifier’s heightened sensitivity to age-related variations in vocal behavior.

The precision of the classifier, gauged at 0.9926, reflected a low false-positive rate, pivotal for precise stress vocalization identification. Simultaneously, the recall rate stood at 0.9936, emphasizing the model’s comprehensive detection of relevant vocal instances across different ages. The harmonized precision and recall culminated in an F1 score of 0.9930, signifying the balanced accuracy and thoroughness of the classifier.

This blend of high precision and recall highlights the classifier’s adeptness in differentiating between the vocal patterns of younger and older hens, a capability crucial for nuanced welfare assessment in poultry farming. Such classification enables a more informed approach to managing and improving the living conditions of laying hens, aligning with the objectives of precision livestock farming and sustainable animal welfare practices.

### 3.16. Changes in Vocalization Patterns Over Time

Further analysis investigated the temporal aspect, exploring how vocalization patterns evolved as the chickens aged. The CNN’s ability to track vocal changes over time provided insights into the developmental vocal adaptations in response to stress. The model highlighted that vocalization patterns are not static; they undergo significant alterations as hens mature, reflecting changes in their behavioral and physiological states in reaction to stressors. The overall results emphasize the intricate relationship between stressor type, age, and vocal responses in laying hens.

### 3.17. Impact of Stressor Timing on Vocalization Patterns

The data reveals a clear shift in vocalization patterns following the application of stressors. The prestress and post-stress vocalizations of the laying hens exhibit distinct profiles, as indicated by the confusion matrix. Despite some overlap in the classification of samples, the model delineates the vocal changes with a reasonable degree of accuracy. This change suggests that the hens’ vocal expressions are immediately influenced by the stress events, and this influence is detectable and classifiable through machine learning techniques.

### 3.18. Vocalization Variations Between Control and Treatment Groups

Comparing the control and treatment groups, the classification results demonstrate discernible differences in vocalization patterns within the same age week. While the acoustic differences may not be apparent to the human ear, the CNN model picks up on subtleties that distinguish the vocal responses of the control group from those subjected to stressors. Over time, the control group’s vocalizations appear to deviate from those of the treatment group, with the five-week control results showing more significant differences than the four-week results. These findings confirms that the vocal patterns of hens are sensitive to environmental conditions and stressors, and these changes are captured in the audio data over time.

### 3.19. Consistency of Vocal Changes Across Treatment Groups and Time

When examining vocal changes over time between ’Treatment 1’ (umbrella stressor) and ’Treatment 2’ (dog barking stressor), the classification model did not distinguish the patterns as distinctly as expected. However, the analysis shows that both treatments induce consistent changes in vocal patterns across different weeks. This consistency indicates that the stressors have a lasting effect on the hens’ vocal behavior, which does not diminish over time as one might expect with habituation. These persistent vocal changes suggest that the stressors continue to impact the hens, an important consideration for welfare monitoring and the development of stress mitigation strategies in poultry farming.

### 3.20. Acoustic Features Interrelationships

Figure 6 displays a detailed correlation matrix, offering insight into the linear relationships between pairs of Mel Frequency Cepstral Coefficients (MFCCs) derived from avian vocal samples. Each cell within the matrix illustrates the correlation coefficient, quantifying the strength and directional association between feature pairs.

**Figure 6.**
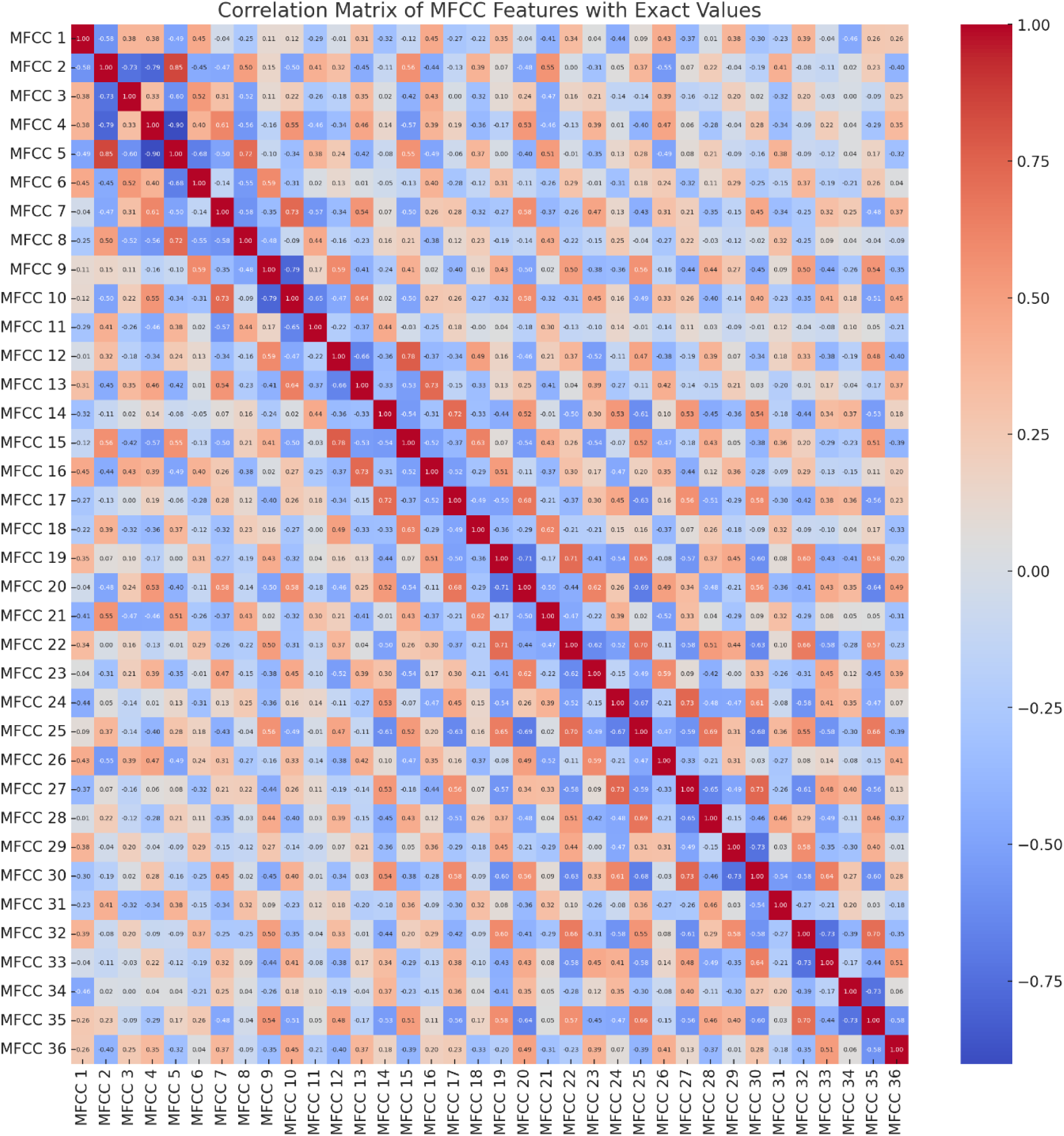
Correlation Heatmap of MFCC Features Across Vocal Samples.

**Figure 7.**
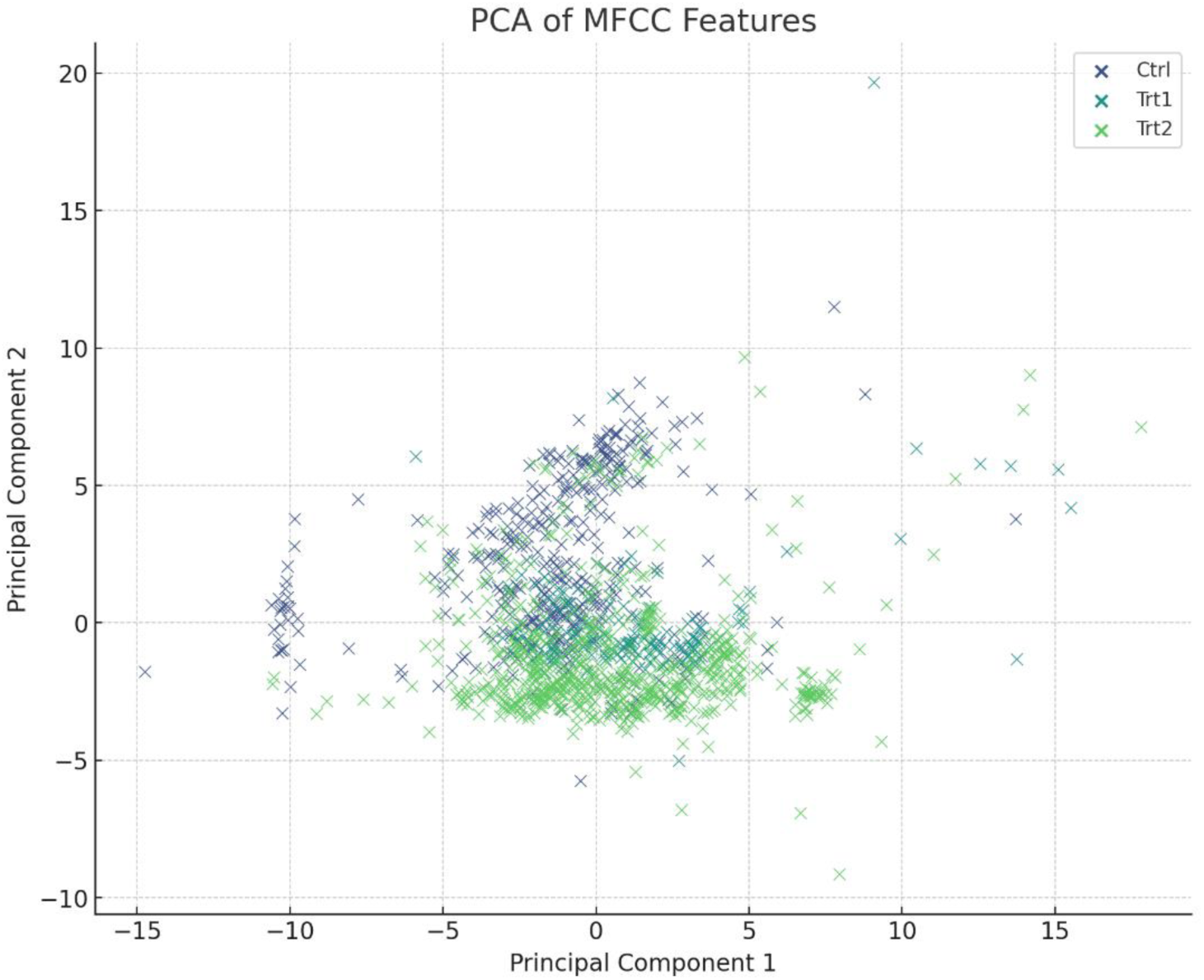
Scatterplot Illustrating the PCA of MFCC Features Across Three Conditions: Control (Ctrl), Treatment 1 (Trt1), and Treatment 2 (Trt2).

#### Strong Positive Correlations

The heatmap’s deep red hues denote cells where correlation coefficients are near +1.00. This coloration highlights MFCC pairs that exhibit synchronous variation, suggesting a parallel increase in their values. Such robust positive correlations indicate that these features potentially encapsulate analogous aspects of the vocalizations. An exemplar of this would be the pair MFCC 1 and MFCC 2, where a hypothetical correlation of +0.90 signifies a potent positive linear relationship, inferring that these features likely respond in concert to vocal modulations.

#### Pronounced Negative Correlations

Conversely, dark blue cells in the heatmap reflect correlation coefficients nearing -1.00. These cells pinpoint MFCC pairs where an uptick in one feature is counterbalanced by a downturn in the other, establishing a strong inverse correlation. For instance, should MFCC 1 and MFCC 3 correlate at -0.85, it would underscore a substantial inverse association, suggesting that these features inversely track the dynamics of the vocal emissions.

#### Mild or Nonexistent Correlations

Cells painted in lighter blue or lighter red shades, with coefficients hovering around zero, characterize MFCC pairs lacking a significant linear relationship. This lack of correlation implies that the features in question contribute unique and independent information to the dataset, potentially encoding distinct vocal attributes. An example might be MFCC 10 and MFCC 20, where a correlation coefficient near zero would indicate no discernible linear dependence, highlighting their individual contributions to the vocal profile.

The heatmap serves as a visual summary of the intricate web of feature interdependencies within the dataset, providing a foundation for informed decisions regarding feature selection in subsequent modeling and analytical endeavors. MFCC features with strong correlations may warrant consolidation or exclusion to preclude information overlap, while those with weak correlations stand out as candidates for investigating the multifaceted nature of avian vocal communication.

The statistical analysis of Mel Frequency Cepstral Coefficients (MFCCs) using Analysis of Variance (ANOVA) yielded significant insights into the vocalization patterns of layer hens under different treatment conditions. The ANOVA results spotlighted the top 10 MFCC features that exhibited the most substantial differences across treatment types, underscoring their potential as key indicators of specific stress responses or developmental changes. Notably, these features demonstrated statistically significant variations, as evidenced by their low P-values, implying that the observed differences were highly unlikely to be due to chance. The most noteworthy features were MFCC 27, MFCC 32, MFCC 8, MFCC 22, MFCC 24, MFCC 21, MFCC 1, MFCC 19, MFCC 16, and MFCC 35, each showing distinct changes in vocalization patterns under varying conditions.

The Principal Component Analysis (PCA) conducted on the Mel Frequency Cepstral Coefficients (MFCC) was depicted in a scatter plot (Figure 7), which demonstrated both distinct separations and notable overlaps among various treatment types. The analysis distilled the MFCC data into two primary components, revealing a pronounced divergence of the control group from the treatment cohorts, thus underscoring the unique influence of treatments on the vocal characteristics. Despite this divergence, overlapping regions suggested the presence of shared vocal feature alterations across the treatments.

The initial two principal components, encapsulating the dataset’s most significant variance, afforded a consolidated overview of these intricate interrelationships. It is important to note, however, that they offer a reductive representation of the complex, multidimensional data. This visualization facilitates an understanding of the broader patterns within the data by reducing its dimensionality. Furthermore, the baseline established from control group data, when compared with pre-stress vocalizations, enabled the identification of intrinsic differences unrelated to stress factors. The PCA not only uncovered patterns indicating how conditions such as control, treatment 1, and treatment 2 affect vocal characteristics but also suggested that including additional features—like pitch, intensity, duration, and formant frequencies—in future studies could provide a more comprehensive picture.

Regarding the graded feature analysis, the investigation of MFCC features revealed changes that evolved gradually over time, potentially reflective of more subtle and enduring effects of stress or aging. By employing scatter plots, the study delineated the differences in features, enabling the identification of both distinct and graded changes in vocalizations. These findings are pivotal in shaping strategies for stress management in animals, offering significant contributions to animal welfare. In synthesizing these observations, we illuminated not only the immediate impact of stressors on animal behavior and communication but also the nuanced, longer-term effects. Such insights are of immeasurable value in the fields of ethology, veterinary sciences, and on-farm animal management.

### 3.21. Impact of Stressor Type on Vocalization

The examination of vocalization patterns in laying hens subjected to distinct stressors, namely the visual stimulus of an umbrella opening and the auditory stimulus of dog barking, sheds light on the intricate sensory processing and adaptive behaviors of chickens. This research emphasizes the importance of discerning these vocal reactions for enhanced poultry welfare monitoring and management. Our findings reveal that chickens’ vocal behaviors are significantly influenced by the nature of the stressor. When exposed to the visual stressor of an umbrella opening, chickens exhibited abrupt and brief vocal responses, indicative of an immediate startled reaction. In contrast, the auditory stressor of dog barking elicited more prolonged vocalizations, suggesting a longer-lasting state of alertness. This variation in vocal patterns demonstrates the chickens’ ability to differentiate between visual and auditory threats, tailoring their vocal expressions to the specific type of stimulus encountered.

#### 3.21.1. Insights from MFCC Feature Analysis

The Mel Frequency Cepstral Coefficients (MFCC) feature analysis played a pivotal role in understanding these vocal dynamics. The heatmap analysis revealed significant variability in the lower-order MFCCs, particularly MFCC 1 and 2. This variability correlates with the chickens’ sharp vocal reaction to the visual stimulus and their more drawn-out response to the auditory stimulus. These lower MFCCs are reflective of major shifts in the audio signal’s energy and spectrum, triggered by the respective stressors.

Further insight is gained from the box plot analysis, where MFCC 1 stands out with a wide range of values and noticeable outliers. This points to a substantial impact of the stressors on the overall energy in the chickens’ vocalizations. On the other hand, higher-order MFCCs exhibit less variability, suggesting they capture more stable and nuanced aspects of the vocal responses to stress.

The scatter plot analysis uncovers complex correlations between pairs of MFCCs, indicating multifaceted vocal responses to different stressors. This intricate correlation structure highlights the chickens’ sophisticated sensory perception system, capable of discerning and reacting variably to diverse environmental cues.

#### 3.21.2. Classification and Practical Implications

Utilizing a Convolutional Neural Network (CNN) model, the study effectively classified the specific acoustic signatures associated with each type of stressor. This classification not only corroborates the distinct vocal patterns in reaction to visual versus auditory stimuli but also serves as a crucial tool for interpreting chicken vocal cues in welfare monitoring scenarios. The analysis of vocalization patterns in response to different stressors offers critical insights into the sensory processing and behavioral adaptation of chickens. It underscores the significance of understanding these vocal responses for developing targeted welfare strategies and informs enhanced management practices in poultry farming.

## 4. Discussion

### 4.1. Detailed Analysis of Vocal Response to Different Stressors

This study has made a significant contribution to our understanding of how laying hens vocally respond to varied environmental stressors. The utilization of Convolutional Neural Networks (CNN) and Mel Frequency Cepstral Coefficients (MFCCs) enabled the discernment of intricate patterns in the vocalizations, revealing how hens distinctly react to different stimuli. For instance, the vocal responses to visual stressors, like an abrupt umbrella opening, contrasted sharply with responses to auditory stressors, such as sudden dog barking sounds. This distinction in vocal patterns underscores the hens’ complex cognitive and emotional processing abilities, highlighting their capacity to differentiate and adaptively respond to various types of environmental changes [44–45]. Such nuanced understanding aids in tailoring stress mitigation strategies and enhances the overall approach to managing poultry welfare.

### 4.2. Age-Related Variations in Vocal Response

The influence of age on vocal response emerged as a pivotal aspect of this research. Significantly, younger chickens displayed vocal patterns that were markedly different from those of older chickens. This variation suggests a developmental trajectory in how chickens perceive and respond to stress [46–47], with younger birds potentially being more sensitive or reactive to environmental changes. Understanding these age-related differences is crucial for developing age-appropriate welfare and management strategies in poultry farming, ensuring that the specific needs of chickens at different stages of their life are effectively addressed.

### 4.3. Immediate and Long-term Impact of Stress on Vocal Behavior

The research demonstrated that stress induction leads to immediate and significant alterations in chicken vocal behavior. This immediate response to stressors is a vital indicator of the birds’ well-being and can be leveraged for early detection of distress [48–50]. Moreover, the persistence of these vocal changes over time points to the potential long-term impacts of stress exposure on chickens, an essential consideration for understanding the cumulative effects of environmental stressors [13] on poultry welfare.

### 4.4. Insights from Vocal Response Analysis to Diverse Stressors

The comparative analysis of vocal responses to different stressors has significant implications for poultry welfare. Recognizing specific vocal responses aids in developing targeted strategies for stress mitigation and informs better management practices. It also highlights the potential of machine learning techniques for refined animal behavior analysis in welfare assessments. This study presents a comprehensive view of chicken vocal behavior in response to diverse sensory stimuli, advancing our understanding of poultry welfare practices. The integration of MFCC feature analysis and machine learning offers a novel approach to deciphering complex vocal responses, underscoring the sophistication of chickens’ sensory perception [9] and their ability to adaptively respond to their environment. The distinction in vocal patterns underscores the hens’ complex cognitive and emotional processing abilities, highlighting their capacity to differentiate and adaptively respond to various types of environmental changes.

### 4.5. Contributions to Poultry Welfare and Precision Livestock Farming

#### 4.5.1. Enhancing Welfare Monitoring Practices

The study’s findings offer transformative potential for poultry welfare monitoring. By non-invasively assessing the well-being of chickens through vocalization analysis, this approach provides a humane and ethical method to detect stress early and respond accordingly [11, 51]. This method aligns with contemporary ethical standards [52] in livestock management, emphasizing the importance of understanding and responding to the animals’ needs proactively.

#### 4.5.2. Advancements in Precision Livestock Farming

The integration of machine learning in analyzing chicken vocalizations represents a significant advancement towards smart farming practices. Employing CNN models and MFCCs for continuous, noninvasive monitoring could revolutionize poultry farming. This technology enables real-time welfare assessments [53–55], leading to more informed decision-making and proactive management strategies, significantly enhancing the efficiency and humaneness of poultry farming.

### 4.6. Future Directions and Applications

#### 4.6.1. Longitudinal Studies on Chronic Stress

Future research should focus on the long-term effects of chronic stressors on hen vocalizations. Investigating whether chickens exhibit habituation or increased sensitivity over time is crucial for understanding their long-term welfare and resilience. Such studies could provide deeper insights into the adaptive capacities of chickens and inform long-term welfare strategies.

#### 4.6.2. Broadening the Scope in Animal Welfare Science

The methodologies used in this research have broad applicability in animal welfare science. Extending these techniques to other species and farming contexts could revolutionize welfare assessments across various animal husbandry domains. This approach promises a more ethical and sustainable future for animal farming, emphasizing welfare-oriented practices. This study has significantly enriched our understanding of chicken vocal behavior under stress and set the stage for future research and applications in poultry welfare monitoring. The integration of advanced technology in animal welfare research embodies a commitment to practices that are not only productive but also ethically responsible and humane. This research paves the way for a future in which animal welfare is not just a consideration but a fundamental aspect of livestock management and care.

## 5. Conclusions

### 5.1. Deciphering Stress-Induced Vocalizations in Poultry - Implications for Welfare Monitoring

The intricate vocalizations of laying hens serve as a communicative bridge, reflecting their internal states and welfare conditions. The comprehensive study conducted herein has systematically unraveled the vocal patterns elicited by environmental stressors, harnessing the analytical prowess of deep learning and the nuanced audio feature extraction capabilities of Mel Frequency Cepstral Coefficients (MFCCs). This fusion of advanced computational methods and acoustic analysis represents a significant stride forward in the endeavor to interpret the subtle lexicon of chicken vocalizations.

Our findings underscore the discernible impact of stressors on vocal expression, varying not only with the nature of the stressor but also with the age of the birds. Younger hens demonstrated vocal patterns markedly distinct from their older counterparts, pointing to a developmental trajectory in their stress response mechanisms. This age-related divergence in vocal behavior, captured through the lens of a Convolutional Neural Network (CNN), highlights the dynamic nature of animal communication and its sensitivity to physiological and environmental stimuli.

The study’s revelations extend beyond the scientific understanding of avian communication. They pave the way for pragmatic applications in poultry farming, particularly in the realm of welfare monitoring. The precision with which the CNN model classified stress-specific vocalizations present a compelling case for the integration of such models into non-invasive monitoring systems. Such systems could potentially detect early signs of distress, enabling timely interventions that mitigate stress and enhance the well-being of the flock.

The correlation between vocal changes and stressors, coupled with the consistent recognition accuracy across various ages and stress conditions, has far-reaching implications. It suggests that continuous monitoring of vocalizations could serve as an effective barometer for the welfare status of poultry. Furthermore, it underlines the importance of tailoring management practices to the nuanced needs of laying hens at different life stages, fostering an environment conducive to their health and productivity. Our research also lays a foundation for future studies to explore the longitudinal effects of chronic stressors on vocalizations and whether hens exhibit habituation or heightened sensitivity over time. Such investigations could offer deeper insights into the resilience of poultry and the long-term implications of their rearing conditions.

In the broader scope of animal welfare science, this study illustrates the utility of bioacoustics and machine learning as powerful tools for advancing our comprehension of animal behavior. The methodology employed here can be extrapolated to other species and contexts, potentially revolutionizing welfare assessments across various domains of animal husbandry.

The integration of CNN models and MFCCs for analyzing chicken vocalizations has provided a novel perspective on the interplay between environmental stressors and animal communication. The implications of this research are manifold, offering a beacon for the development of enhanced welfare monitoring systems and informing best practices in poultry management. Ultimately, this study serves as an affirmation of the potential that lies in the intersection of technology and animal welfare to foster practices that are not only productive but also ethically grounded and humane.

### Funding

This research received no external funding.

### Institutional Review Board Statement

The study was conducted in accordance with the Declaration of Helsinki, and approved by the Institutional Review Board (or Ethics Committee) of the Wageningen University & Research, The Netherlands.

### Data Availability Statement

The data supporting the findings of this study are openly available in the Zenodo repository. This includes five figures integral to the research, an Excel file containing the dataset of 40 Mel Frequency Cepstral Coefficients (MFCC) features, and processed audio data of laying hen vocalizations. While the processed data set is approximately 2.5 in size, the original raw data, amounting to about 8 GB, is not included in the repository but can be made available upon reasonable request. Additionally, the repository provides access to Python algorithms used for CNN feature extraction and classification in the study, as well as files detailing the classification results and confusion matrices. Interested parties are encouraged to access these resources via the link https://doi.org/10.5281/zenodo.10433023 to explore the comprehensive dataset and methodology underpinning this research.

## Supporting information

Supplemental 1 - MFCC Features

## Acknowledgments

This research was greatly enriched by data collected during a comprehensive collaborative project at the Animal Sciences Department, Wageningen University & Research. Profound appreciation is extended to the Adaptation Physiology Group, with special thanks to Naomi Pijpers, Huib van den Heuvel, and Ali Youssef, who worked diligently under the guidance of Dr. Suresh Neethirajan. Their meticulous efforts in data collection were pivotal to the success of this study. It is, however, crucial to acknowledge that while the data collection was a collaborative effort, the conceptualization, development of the algorithm, analysis of the data, and the drafting of this manuscript were independently executed by Dr. Neethirajan. His expertise and vision have been instrumental in the realization of this research, providing critical insights and direction throughout the process. The depth and breadth of the analysis presented in this study are a testament to his significant contributions to the field of poultry vocalization research.

## Conflicts of Interest

The authors declare no conflicts of interest.

## Notes

https://doi.org/10.5281/zenodo.10433023

### Competing Interest Statement

The authors have declared no competing interest.

### Summary of Updates

The revised manuscript now includes results from further data analysis.

